# Impact of Glycan Depletion, Glycan Debranching and Increased Glycan Charge on HIV-1 Neutralization Sensitivity and Immunogenicity

**DOI:** 10.1101/2024.02.20.581329

**Authors:** Alessio D’Addabbo, Tommy Tong, Emma T. Crooks, Keiko Osawa, Jiamin Xu, Alyssa Thomas, Joel D. Allen, Max Crispin, James M. Binley

## Abstract

Modifying HIV-1’s envelope glycoprotein glycans can impact its neutralization sensitivity. The use of the knock out cell line GnT1-prevents the elaboration of complex-type glycans, and opens up the glycan shield, increasing bNAb vulnerability. Some bNAb precursors can bind to GnT1-trimers, supporting their use in vaccine priming. However, GnT1-trimers express poorly and exhibit very low infectious counts. Here, we describe two other potentially vaccine-relevant glycoengineered trimers, 1) To truncate complex glycans, we used of a cocktail of glycosidases, termed “NGAF3” (<u>N</u>euraminidase, β-<u>G</u>alactosidase, N-<u>A</u>cetylglucosaminidase and endoglycosidase <u>F3</u>). Like GnT1-trimers, NGAF3 reduced glycan clashes and increased bNAb potency while retaining a closed trimer conformation; 2) Modified by β-1,4-galactosyltransferase 1 (B4GalT1) and β-galactoside α-2,6 sialyltransferase 1 (ST6Gal1) during Env biosynthesis. Glycan mass spectrometry revealed that NGAF3 removed glycan heads of 3 of 7 positions on the trimer that are largely occupied by complex glycans. It also revealed a novel B4GalT1 activity to favor glycan precursor conversion to hybrid glycans rather than complex glycans. A comparison to monomeric gp120 revealed that B4GalT1’s new activity depends on tight glycan spacing. B4GalT1 affected more glycans than NGAF3 (6 out of 7 glycans), perhaps due to greater accessibility *during* Env folding rather than *after* folding. Surprisingly, the N611 glycan was unaffected by either modification. B4GalT1 and ST6Gal1 cooperatively increased the abundance of hybrid glycans and α-2,6 hypersialylated termini. This occurred largely by amplifying the abundance of a limited number of hybrid glycan structures that are also present on unmodified trimers. In rabbit vaccinations, B4GalT1+ST6GalT1-modified virus-like particles reduced the frequency and titers of serum NAbs that showed a modest preference for modified glycans. Conversely, chronically HIV-1-infected donor plasma neutralizing antibody titers were 1.7- to 10.8- fold higher against B4GalT1+ST6GalT1-modified pseudovirus. Overall, our data provide tools for heterologous prime, boost and polishing vaccine regimens using modified glycans.

**AUTHOR SUMMARY:** An HIV-1 vaccine remains one of the most significant biomedical challenges today. Vaccines often work by triggering virus fighting antibodies to stave off infection. For HIV-1, this has been exceedingly difficult because the target, called the HIV-1 envelope glycoprotein (Env) is extremely variable and carries a thick sugar coat, protecting it from all but a few rare broadly reactive antibodies (termed bNAbs) that sometimes develop during natural HIV-1 infection. Given these challenges, it is reasonable to propose that any successful vaccine will need to be rationally designed to trigger the rare bNAbs. To meet this goal might require more 3 different vaccine components. First, a vaccine “prime” to trigger rare bNAb proliferation, followed by heterologous “boosts” and “polishing” immunogens to select for bNAbs. In this study, we evaluated two potential immunogens. In one approach, we used cocktails of enzymes to strip Env’s sugar coat, in the hope of reducing the barrier to stimulate bNAb precursors. In another, we used excess enzymes to build particular structures on Env’s coat. We checked how well these two Env variants were recognized by antibodies. We also checked in fine detail how the sugars were changed at the molecular level. Finally, we immunized rabbits. Our data enrich the number of strategies available to further explore the concept of priming, boosting and polishing vaccine shots.

## INTRODUCTION

Immune tolerance dictates that antibodies (Abs) generally avoid glycans of viral glycoproteins derived from human cells [1]. HIV-1 Env trimer glycosylation density is exceptionally high, comprising approximately half of its mass. This carbohydrate shell leaves few glycan-free holes of sufficient size to admit broadly neutralizing antibodies (bNAbs). As a result, bNAbs usually take at least a year to develop in about 20% of infected people [2]. The scores of new bNAbs recovered from HIV-1 infected donors since 2009 [3] suggest that despite their preference for protein epitopes, bNAbs inevitably develop some glycan contacts and avoid clashes with others [4–6].

NAb breadth can be blunted by strain-specific glycans. For example, V2-specific bNAb neutralization is constrained by neighboring glycans at N130 and/or the C-terminal portion of the V2 loop [5, 7]. Some sequons are incompletely occupied in soluble Env vaccine trimers, resulting in non-encoded “glycan holes”. By comparison, “sequon skipping” is absent in native, full-length trimers, making them invulnerable to Abs that recognize these non-encoded holes [8, 9].

Maximum percent bNAb neutralization (i.e., bNAb saturation) may be impacted by glycan heterogeneity. Thus, a bNAb may neutralize a fraction of virus with a relatively small clashing glycan, but not the fraction with a large clashing glycan [10, 11]. This epigenetic virus “escape” results in a sub-saturating neutralization plateau [5, 10–13]. Because of these challenges, preclinical vaccine NAbs tend to vary in potency and saturation, usually with limited breadth [14–17]. Notably, increasing the size of the glycan hole at the CD4 receptor binding site by depleting the glycan fence improves NAb induction against pseudoviruses (PVs) that lack these glycans, but not against the parent [15, 17–19].

We previously investigated the effects of 16 different glycoengineering methods on HIV-1 NAb breadth, potency and saturation [5]. This revealed that some bNAb ancestors bind more effectively to and/or neutralize GnT1-modified PV, in which large complex glycans are replaced by smaller Man5 high mannose glycan (Fig. 1B) [5, 20]. In some cases, mature bNAbs continue to remain most potent against GnT1-PV (e.g., CH01, 35O22, VRC34), as clashes with other glycans limit their potency and/or saturation. Indeed, some but not all HIV-1+ donor plasmas also show increased NAb potency against GnT1-PV [5]. Since GnT1-trimers retain a “closed” conformation [5], they might be of particular use to trigger bNAb precursors [5, 20]. The use of GnT1-cells is limited by ∼10-fold reduced Env expression and usually markedly reduced PV infectivity counts [21]. In the current study, we attempted to overcome this expression/infectivity problem by treating PV with glycosidases *ex vivo* to deplete complex glycans and relax the constraints on NAb binding (Fig. 1C).

**Fig 1.**
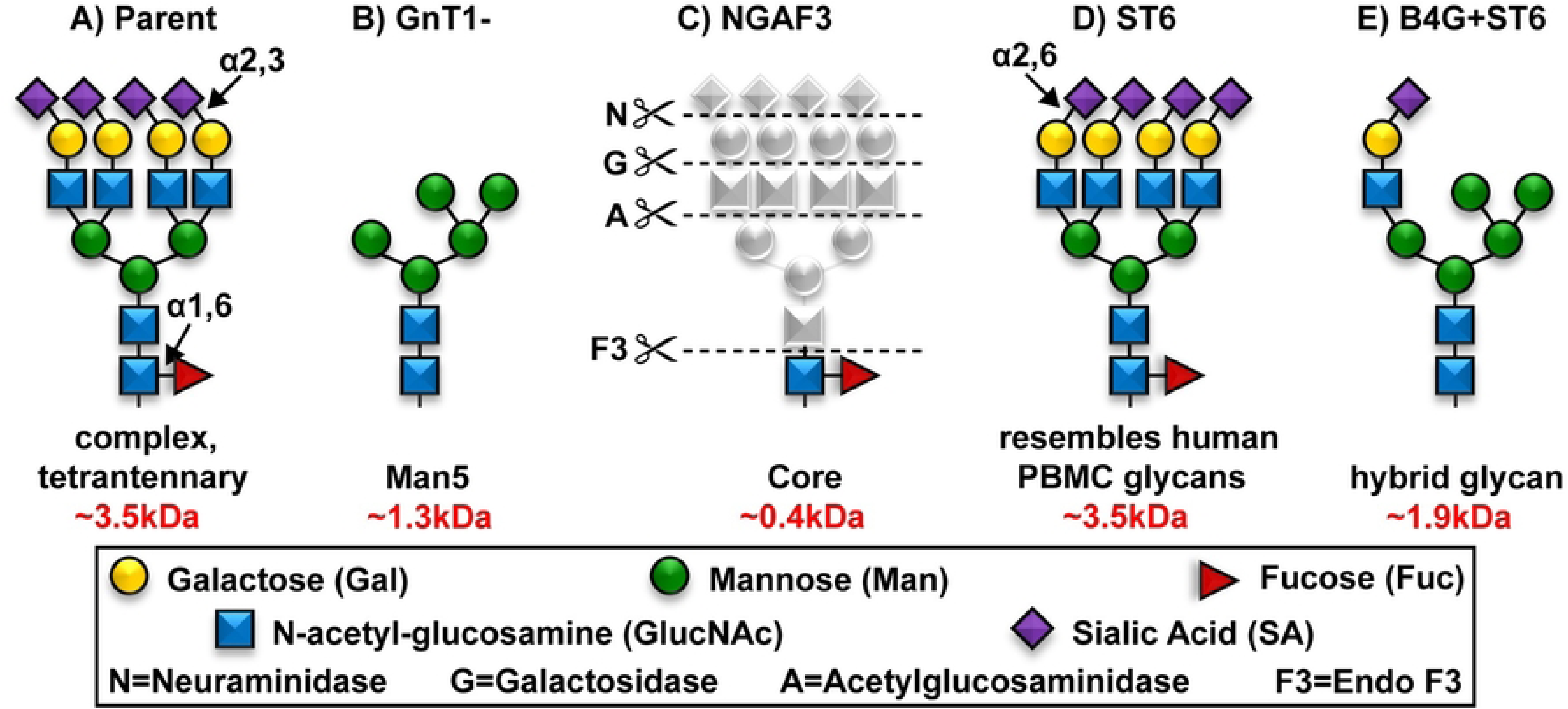
Schematic of glycan modification methods. (A) Complex glycan, as found in 293T-expressed Env, bearing terminal α-2,3 sialic acid. (B) Man5 glycan, as produced in GnT1-cells. (C) NGAF3 digestion by a glycosidase cocktail (abbreviation in parentheses) of neuraminidase (N), galactosidase (G), acetylglucosaminidase (A) and endo F3 (F3) reduces complex glycan to core. Endo F3 specifically cleaves fucosylated biantennary glycans and afucosylated triantennary glycans. (D) A complex glycan produced with co-transfected plasmid expressing ST6Gal1 transferase (ST6) that promotes α-2,6 sialic acid termini, resembling human PBMC-produced HIV-1. (E) Hybrid glycan produced by co-transfecting Env with B4GalT1 transferase (B4G) and ST6. Molecular weights of each glycan type are indicated.

Although bNAbs may benefit from reduced glycan clashes, in many cases they also develop glycan head group contacts [4, 5]. Notably, certain V2 bNAbs (e.g., CAP256, PG9) require terminal α-2,6 sialic acids for optimal neutralizing activity [22]. PVs produced in 293T cells exhibit largely α-2,3 sialic acid termini (Fig. 1A), while peripheral blood mononuclear cells (PBMC)-produced HIV-1 is hypersialylated with α-2,6 sialic acid termini (Fig. 1D) [5, 23–25]. As a result, co-transfection with β-galactoside α-2,6-sialyltransferase 1 (ST6Gal1, henceforth abbreviated as ST6) greatly improves PV sensitivity to V2 bNAbs CAP256 and PG9 [5]. Infectious molecular clones (IMCs) grown in PBMCs also show improved sensitivity to these bNAbs compared to IMCs produced in 293T cells [5]. Thus, it may be important to make vaccines with glycan profiles that resemble those on natural HIV-1 virions.

SDS-PAGE-Western blot analysis revealed that co-transfection of Env with the glycosyltransferase B4GalT1, henceforth abbreviated to B4G, increases gp41 endo H sensitivity, suggesting that complex glycans are replaced with hybrid glycans [5]. We inferred that B4G diverts glycoprotein traffic towards elongation, thereby dampening lateral glycan branching and replacing them with hybrid glycans (Fig. 1E). If so, this would be an unprecedented B4G activity, which is known only to transfer galactose onto N-acetylglucosamine substrates. To test if this new putative activity depends on the unusually close glycan packing, we compared B4G effects on glycan maturation of HIV-1 viral Env, monomeric gp120 and a representative monoclonal antibody (MAb) by glycopeptide mass spectrometry. Lastly, we immunized rabbits with VLPs bearing Env trimers with or without B4G+ST6 modification to see the combined effects of glycan thinning and hypersialylation on Ab responses. Our results provide molecular clarity on the effects of these modifications on HIV-1 native trimer glycans, their neutralization sensitivity and immunogenicity.

## RESULTS

Given the apparently limited glycoprotein expression capacity of GnT1-cells, we sought an alternative way to reduce native trimer glycan clashes. Considering glycosidase digestion, one option is endoglycosidase H (endo H) that partially removes glycans but is sterically limited by high glycan density [26, 27]. Given the greater accessibility of complex glycans to glycotransferases during glycoprotein folding, they might be expected to be more sensitive to *ex vivo* glycosidase removal. PNGase F cleaves at the base of complex glycans. However, it deamidates the glycan-linked asparagine to aspartic acid. Moreover, since glycans tend to cover hydrophobic patches, their complete removal may disrupt protein structure. Conversely, truncating glycans may be a good compromise to retain some coverage of hydrophobic protein patches. Recently, we showed that neuraminidase can remove sialic acid termini from Env trimers [5]. A glycosidase mixture might collectively remove further glycan subunits. We used neuraminidase (abbreviated as ’N’), galactosidase (’G’), acetylglucosaminidase (’A’) in cocktails with or without endo F1, F2 and/or F3 (abbreviated as F1, F2 and F3) to digest JR-FL E168K+N189A PVs, our prototype strain carrying mutations to knock in V2 bNAb sensitivity. Endo F1, F2 and F3 digest native proteins between the diacetylchitobiose core of the glycan stem (Fig. 1C). Endo F1 preferentially digests hybrid and oligomannose structures; endo F2 preferentially digests biantennary glycans and endo F3 preferentially digests fucosylated bi- and afucosylated tri-antennary complex glycans. We compared the effects of these glycosidases in various combinations on PV neutralization.

Digestion by the full NGAF123 cocktail increased PV sensitivity to V3 MAb 39F (Fig. 2B) and reduced PV sensitivities to V2 NAbs PGT145, PG9 and CH01 (Fig. 2D-2F), suggesting an “open” tier 1-like trimer conformation. All digests decreased PGT151 sensitivity, suggesting galactose depletion (Fig. 2H) [5]. Conversely, the glycan fence surrounding VRC01 (Fig. 2A) and the high mannose patch (PGT121; Fig. 2C) were unaffected [15]. CH01 sensitivity was increased by all treatments that included NGA, except for NGAF123 (Fig. 2F). Therefore, depleting glycans at the N-acetyl glucosamine structures is sufficient for maximum CH01 neutralization, as long as a closed conformation is retained (Fig. 1C). PGT145 sensitivity also moderately increased with digests (except NGAF123) (Fig. 2D). Interestingly, PG9 sensitivity was slightly reduced by NG treatment, presumably due to the loss of sialic acids that contribute to its binding [5, 28, 29]. However, further digestion by additional enzymes restored some PG9 sensitivity, presumably by resolving minor clashes. Sub-saturating 35O22 neutralization was also partially resolved by NGAF3 and NGAF23 (Fig. 2G; maximum neutralization saturation of 81% and 80%, respectively, compared to other treatments showing 55-67%). Overall, these glycosidase cocktails markedly impacted bNAb sensitivities. NGAF1 PV sensitivity was like NGA, suggesting that endo F1 had little effect, except in the context of NGAF123. NGAF23 sensitivity was little different than NGAF3. Given the broader specificity of endo F3 and its ability to digest NGA-depleted trimannosyl cores, we selected NGAF3 for further study.

**Fig 2.**
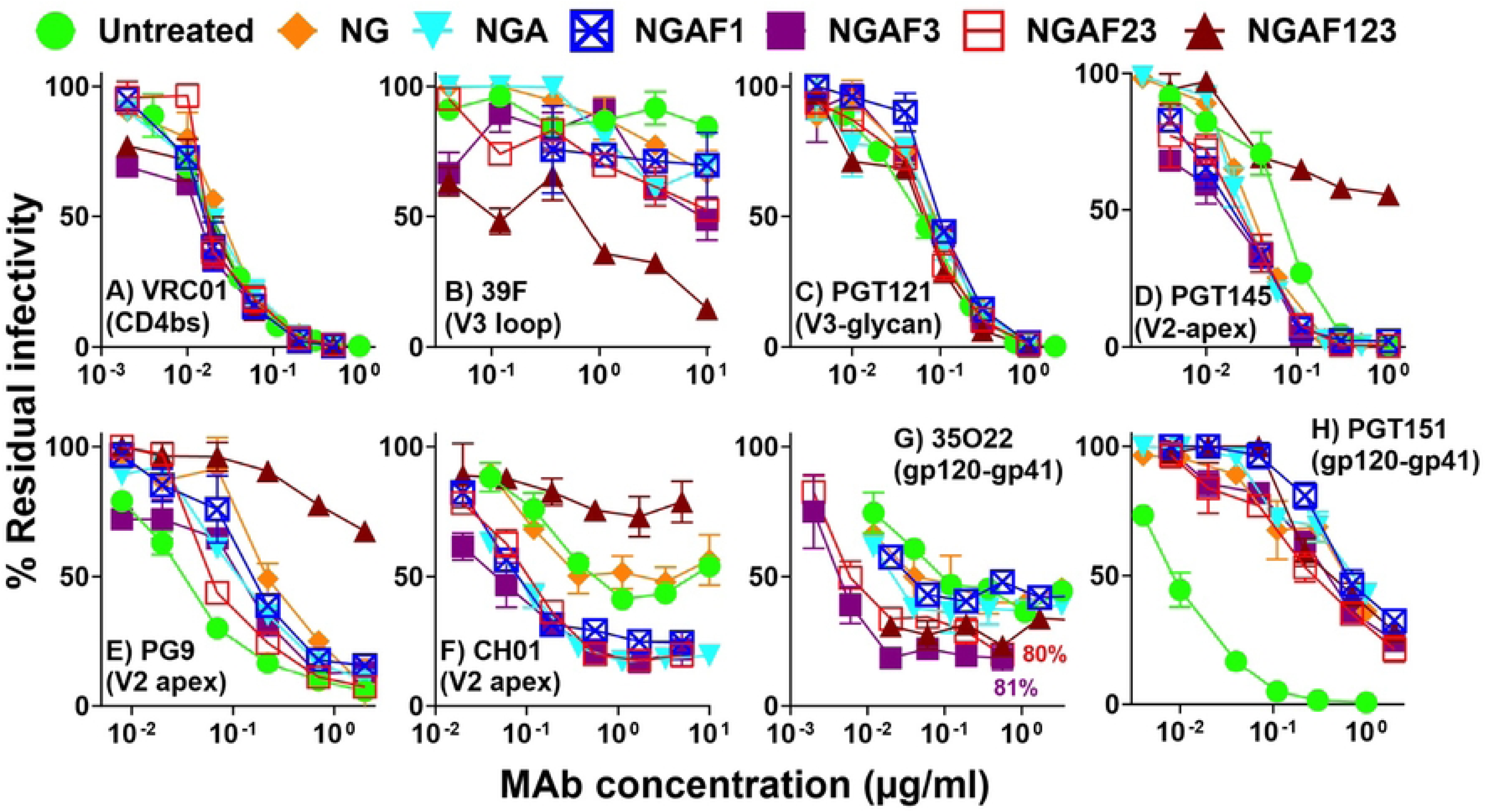
Effects of various glycosidase treatments on pseudovirus neutralizing sensitivity. JR-FL WT gp160ΔCT E168K+N189A PV was treated with various glycosidases for 1h at 37°C. PV sensitivity to various mAbs was assessed by neutralization assay using CF2.CD4.CCR5 cells. Glycosidases are abbreviated as follows: neuraminidase (N), galactosidase (G), acetylglucosaminidase (A), endo F1 (F1), endo F2 (F2), endo F3 (F3). Error bars represent the standard deviation of the mean. Percent maximum neutralization (saturation) is indicated at the highest 35O22 concentration.

Contrasting sharply with poorly infectious GnT1-PV, NGAF3 PV was more infectious even than parent PV (Fig. 3A). ST6- and B4G+ST6-modified PV infectivities were comparable to parent PVs (Fig. 3A) [5]. The A328G mutant was only moderately infectious, as is generally true for “open trimer” mutants (Fig. 3A) [8, 15, 16].

**Fig 3.**
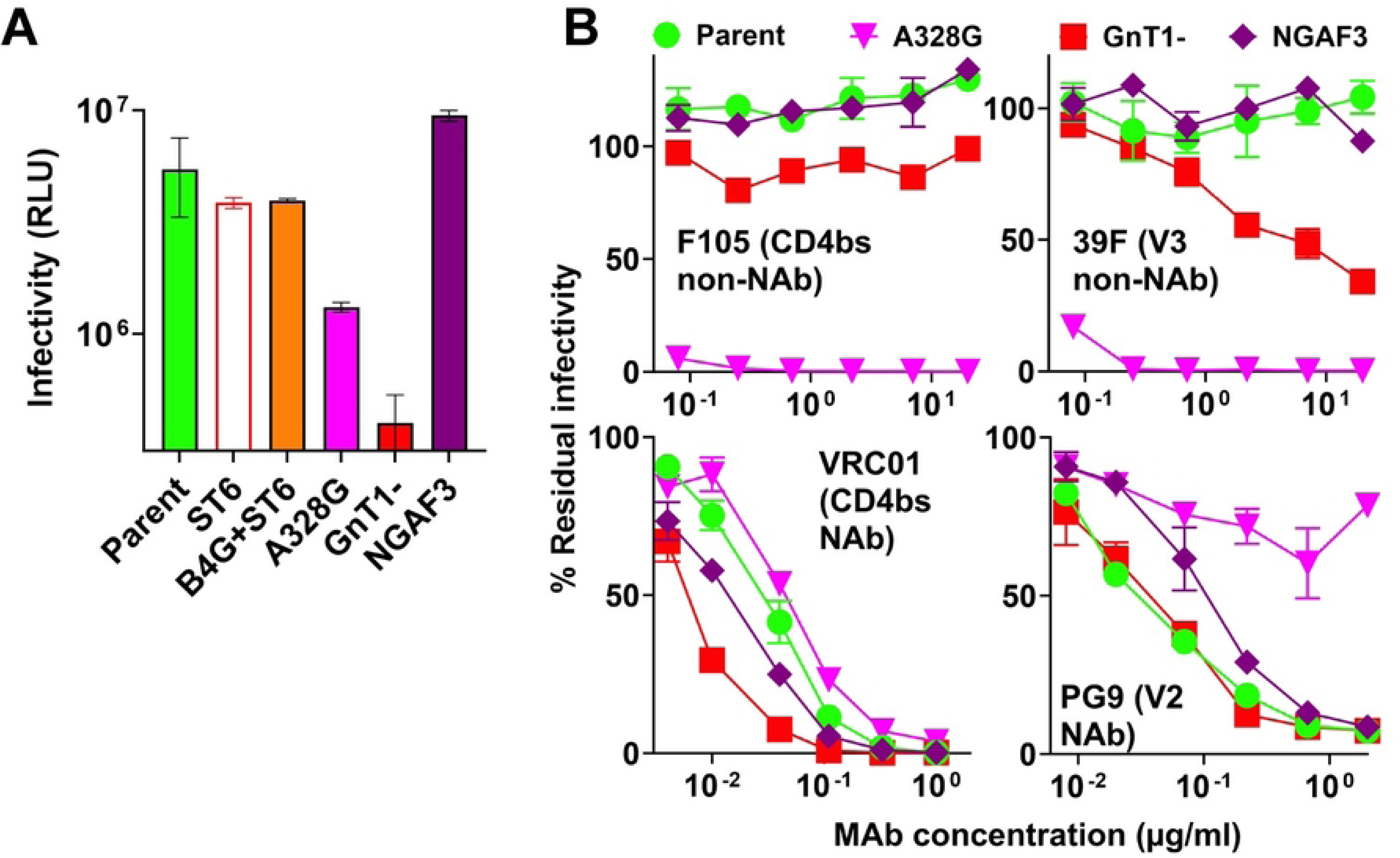
Infectivity and neutralization sensitivity of JR-FL WT gp160ΔCT E168K+N189A PV variants. (A) The infectivity of various JR-FL WT-based PVs on CF2.CD4.CCR5 cells were measured in RLU (relative light units). (B) PV neutralization sensitivity to various mAbs was evaluated using CF2.CD4.CCR5 as target cells. In both assays, the error bars represent the standard deviation of the mean.

We previously described the NAb sensitivity profiles of ST6- and B4G+ST6-modified PVs [5]. In Fig. 3B, we compared NGAF3, A328G, GnT1- and parent PVs. A328G was vulnerable to V3 and CD4bs non-NAbs, but NGAF3 and parent PVs were both resistant. GnT1-PV also resisted F105 but was slightly sensitive to V3 MAb 39F. VRC01 was more potent against GnT1- and NGAF3 PVs. A328G PV was slightly less sensitive than parent PV, perhaps due to its altered conformation. PG9 neutralized parent and GnT1-PV comparably but was less effective against NGAF3 and only marginally effective against A328G (Fig. 3B). Adding gp120 3mut (I423M+N425K+G431E) reduced VRC01 potency against all 4 PVs (S1 Fig). It also interfered with F105 and 39F neutralization of A328G, and 39F neutralization of GnT1-PV. As expected, V3 peptides interfered with 39F neutralization of A328G and GnT1-PVs. Lastly, PG9 was unaffected by either V3 peptides or gp120. Overall, GnT1- and NGAF3 PVs both retained a tier 2 phenotype despite a weaker glycan shield, whereas A328G causes non-NAb-sensitive open trimers.

### How do these modifications impact the V2 sensitivities of other strains?

In Fig. 4A, we tested the effects of B4G+ST6, GnT1- and NGAF3 on the V2 NAb sensitivities of 4 strains. VRC38 and CH01 and their precursors both prefer small glycans. Like GnT1-, NGAF3 increased JR-FL E168K sensitivity to VRC38, VRC38 mHgL, and CH01, but remained CH01 UCA-resistant (Fig. 4A). Also like GnT1-, NGAF3 increased WITO sensitivity to VRC38 and CH01 and their precursors (Fig. 4A). In contrast, the Q23 parent was already sensitive to all 4 MAbs and was not greatly affected by these modifications. GnT1- and NGAF3 both improved c1080 sensitivity to VRC38.01 mHgL, but other MAbs were not greatly affected. B4G+ST6 improved PG16 sensitivity, consistent with its apparent preference for α-2,6 sialylated hybrid glycans [5, 30]. However, it did not consistently improve sensitivity to the other MAbs or their precursors. PGT145 sensitivities of different strains were affected in unpredictable ways by these modifications. 39F did not neutralize any strains regardless of modification. Overall, NGAF3 and GnT1-modifications may have increased potential to engage NAb precursors.

**Fig 4.**
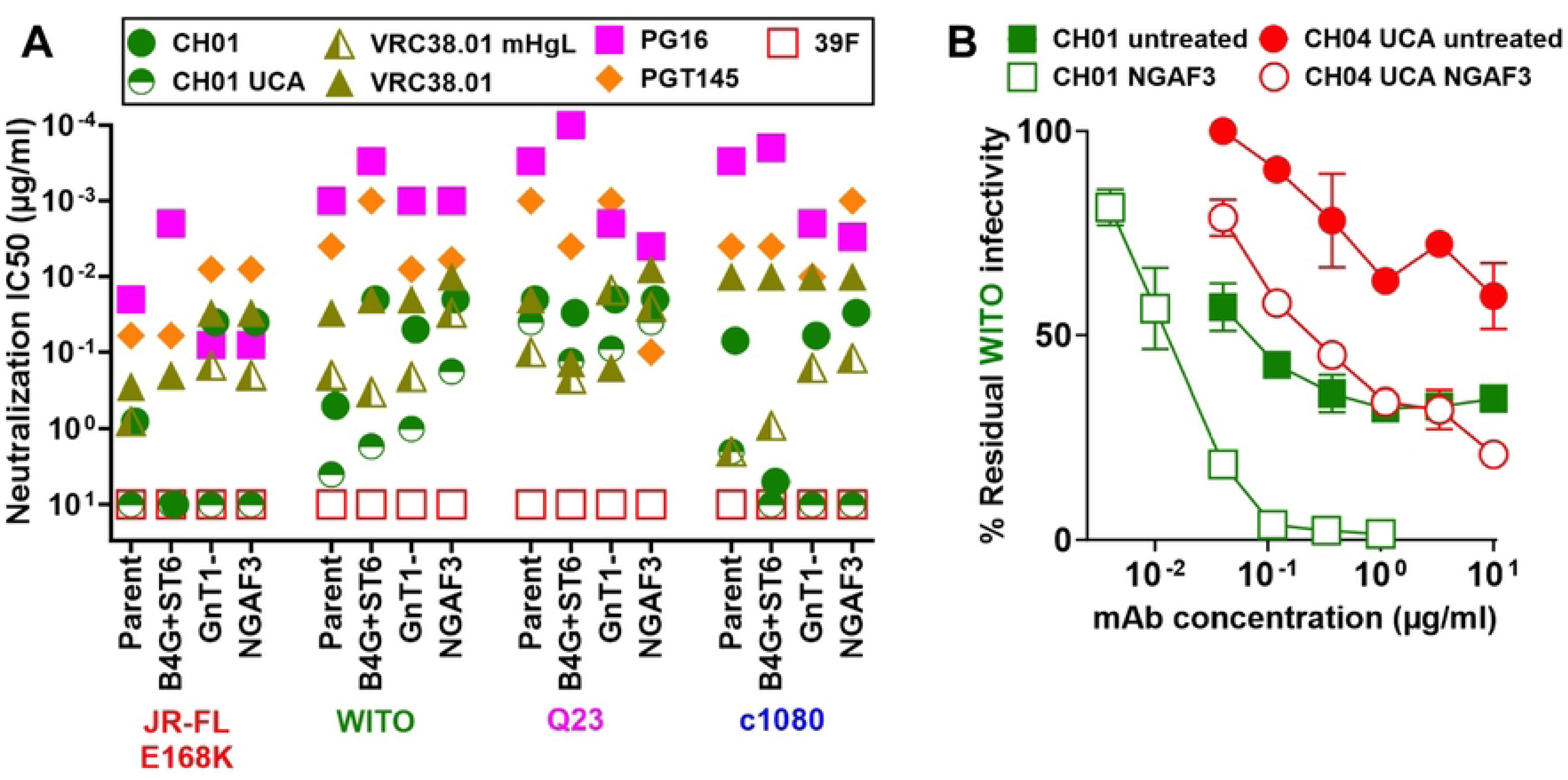
Effects of GE and NGAF3 glycosidase treatment on the NAb sensitivities of different HIV-1 strains. (A) Neutralization sensitivity of JRFL WT gp160ΔCT E168K (subtype B), WITO SOS gp160ΔCT (subtype B), Q23.17 SOS gp160ΔCT (subtype A) and c1080 SOS gp160ΔCT (subtype AE) PVs in parent or modified formats, as shown. (B) CH01 UCA and mature CH01 neutralization of WITO SOS gp160ΔCT with and without NGAF3 treatment. Error bar represents standard deviation of the mean of two independent experiments.

Next, we compared these JR-FL Env variants by blue native PAGE (BN-PAGE). B4G ran slightly faster than the unmodified parent, perhaps due to changes from complex to hybrid glycans (S2A Fig, lane 2). ST6 ran slightly faster (S2A Fig, lane 3), possibly due to hypersialylation, which increases negative charge [5]. In contrast, weakly expressed GnT1-trimers (S2A Fig, lane 4) and NGAF3 trimers (S2A Fig, lane 5) migrated slower than the parent, perhaps due to diminished sialic acid content. The B4G+ST6 trimer mobility was similar to ST6 alone (S2A Fig, compare lanes 3 and 6). In a previous study, A328G did not perceptibly impact Env mobility in BN-PAGE, but exhibited more monomer and less trimer, suggesting decreased lateral stability [8]. Overall, NGAF3 provides an alternative way to reduce complex glycans without the weak expression and infectivity associated with GnT1- (Fig. 3A).

In SDS-PAGE, bands are annotated by colored dots, indicating two gp160 isoforms: ‘mature glycan’ gp160 (gp160m, magenta dots) and immature (high mannose) gp160 (gp160i, yellow dots) (S2B Fig) [8]. Other Env species include mature, functional gp120 (red dots) and gp41 (green dots). ST6 band patterns resembled the parent (S2B Fig, lanes 1-4), as sialic acid changes are unlikely to impact SDS-PAGE (negative charge is saturating). In contrast, with endo H, B4G+ST6 gp120 and gp160m moved faster than the parent (S2B Fig, compare lanes 1 and 2 to lanes 5 and 6). Gp160i was unaffected as this is an early misfolded isoform that is unaffected by modifications [8]. B4G+ST6 gp41 was partially endo H-sensitive, resulting in a ladder effect. A328G was almost identical to the parent (S2B Fig, compare lanes 1, 2, 7 and 8). GnT1-Env lacks complex glycans that impart glyco-heterogeneity, consistent with the diffuse gp160m, gp120 and gp41 bands of the parent. In contrast, the poorly expressed GnT1-gp160m, gp120 and gp41 bands were all sharp, consistent with greater homogeneity (S2B Fig, compare lanes 1 and 2 to lanes 9 and 10). Endo H removed all GnT1-glycans (S2B Fig, compare lanes 9 and 10). Two gp41 bands were observed (S2B Fig, lane 9, α-gp41, lower blot). The lower band may lack glycan at one or more positions. These two bands coalesced into a single fully glycan-depleted gp41 band with endo H (S2B Fig, compare lanes 9 and 10 in α-gp41, lower blot).

We next evaluated the effects of NGAF3 cocktail components. Env treated only with neuraminidase (N) migrated like the parent (S2B Fig, compare lanes 1, 2 to lanes 11 and 12). Mixing neuraminidase and galactosidase (NG) visibly increased gp160m, gp120 and gp41 mobility, which was clearer in lanes digested with endo H (S2B Fig, lanes 11-14). Further mobility changes were not detected when acetylglucosamidase (NGA) and endo F3 (NGAF3) were added (S2B Fig, compare lanes 13-18), suggesting that steric constraints limit their activities. The diffuse nature of NGAF3’s gp160m, gp120 and gp41 bands suggest that some complex (i.e., heterogeneous) glycans survive. Overall, NGAF3 partially reduces the glycan shield, while B4G and ST6 drive hybrid glycans and hypersialylation, respectively (S2A Fig). NGAF3 and B4G efficacies may be limited by the variable glycan density over the trimer surface. Isolated glycans might be more NGAF3-sensitive, while closely packed glycans may be most effectively driven by B4G to become hybrid instead of complex.

### Probing the effects of modifications by glycopeptide mass spectrometry

We used liquid chromatography-mass spectrometry (LC-MS) glycopeptide analysis to determine how modifications impact glycans across the trimer surface on JR-FL SOS gp160ΔCT E168K+N189A parent, B4G+ST6 and NGAF3 VLPs. Glycopeptide data is shown in S1 Table, in which each sheet shows data from different experimental runs. Glycotypes were assigned a score between 1 (Glc1Man9GlcNAc2, hereafter referred to as Man9Glc (glucose)) and 19 (a highly differentiated complex glycan (HexNac(6)(F)(x)) [8]. Each sequon was then assigned a score determined by the sum of each glycotype scores multiplied by its abundance (S1 Table, summarized in S2 Table “Glycan scores” sheet). Averaging parent VLP data in different runs [8, 31] improves its accuracy (S1 Table “Average parent VLP” sheet). It also eliminate gaps in experimental runs in which glycopeptide data is unscorable for technical reasons. Small score variations between the parent samples in different runs may in part be due to the instrument rather than true sample variation. The abundance of various glycan subunits at each site is summarized in each sheet of S2 Table. Sites occupied by predominantly complex glycans are most prone to change. Since glycan scores are based on average data, complex changes may be overlooked. To address this problem, glycan distributions are shown in Fig. 5. Average parent glycan scores are modeled in Fig. 6A, in which high mannose glycans are shown in green and complex glycans are shown in magenta. Glycans that are neither predominantly high mannose nor complex are shown in white and those for which we could not generate data are shown in blue.

**Fig 5.**
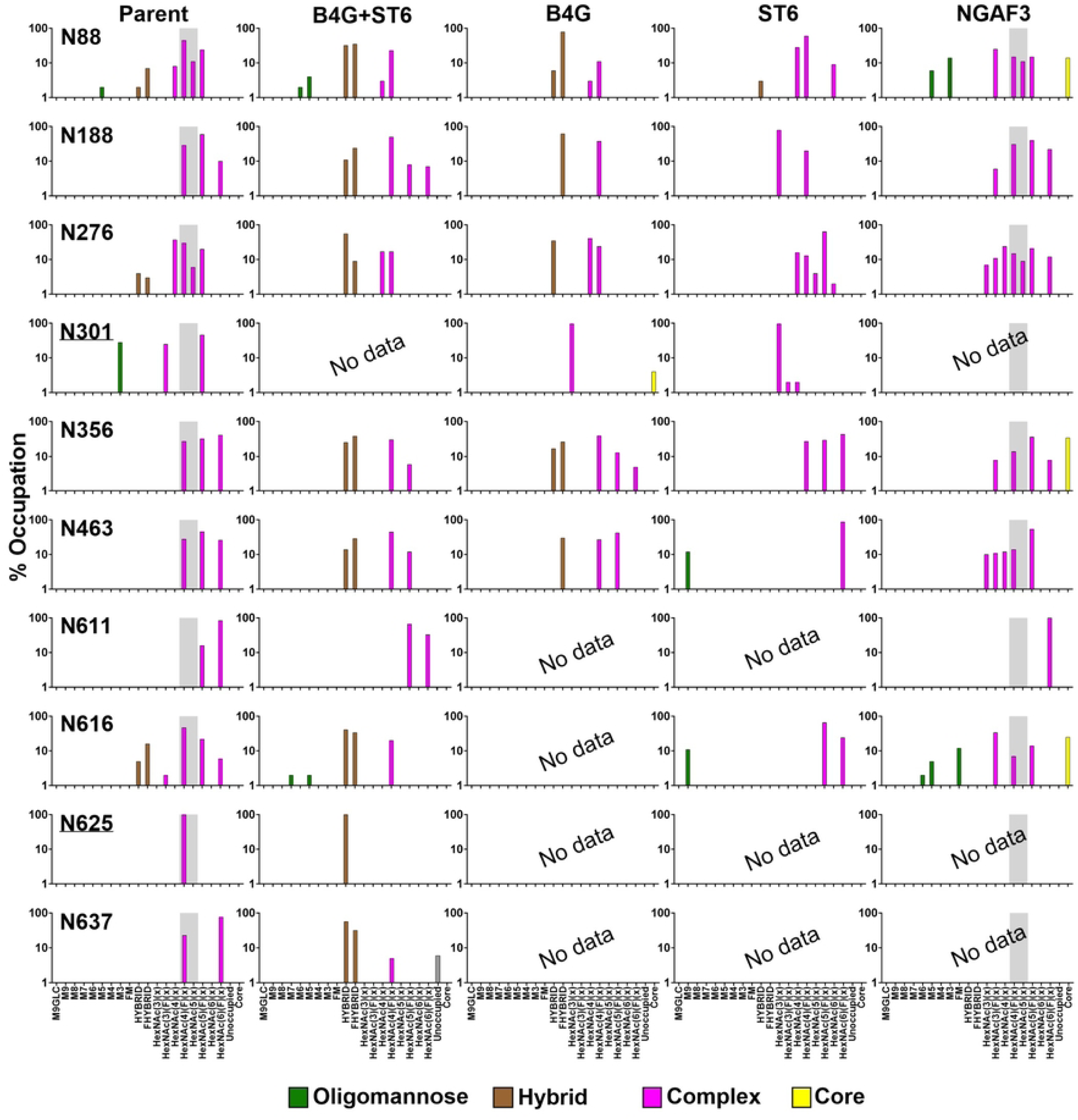
Changes in glycan distribution at key sites of VLP Env. Related to S1 Table and S2 Table. Glycan positions are numbered according to HxB2 strain. Distribution of glycan species is shown at sites predominantly occupied by complex glycans. Gray shading depicts complex glycans that are known to be endo F3-sensitive. At most positions, the matched Parent VLP is compared to B4G+ST6, B4G, ST6 and NGAF3 VLP to show how modifications affect the quantities and types of glycan at each site. For N301 and N625 (underlined), we show the average Parent VLP data because data was missing in the matched Parent VLP. Positions at which no glycopeptide was detected are indicated as “No data”.

**Fig 6.**
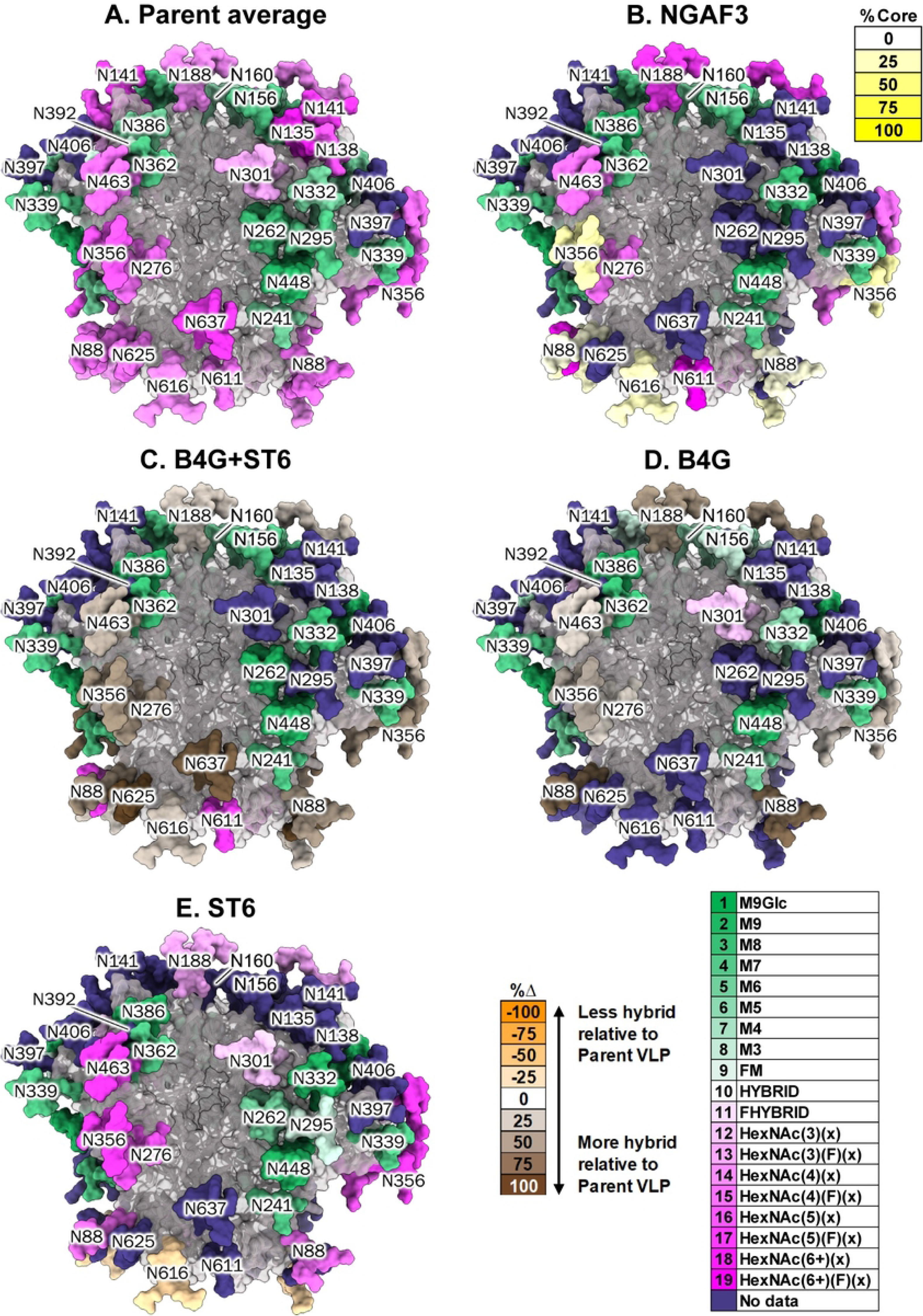
Structural models of trimer glycan changes. Related to S1 Table and S2 Table. (A) Parent VLP (average) glycan scores are assigned on a trimer model (PDB: 6MYY). Each glycan is labeled with its canonical number. Complex glycans are shown in shades of magenta. Oligomannose glycans are shown in shades of green. Glycans for which there was no data are colored blue. Glycan difference (expressed as %Δ) of the following VLPs are compared with parent VLP. Large % differences between VLPs are indicated by progressively darker color. (B) NGAF3-digested trimer model. Complex glycans that became substantially core are shown in shades of yellow. Unaffected glycans are shown with their colors as in part A. (C) B4G+ST6, (D) B4G, and (E) ST6 trimer models. For parts C - E, complex glycans that become substantially hybrid are shown in shades of brown. Unaffected glycans are shown with their colors as in part A.

### i) Effects of NGAF3 on VLP Env

Above, NGAF3 improved NAb binding (Figs. 2-4), although complex antennae appeared to be only partially removed. NGAF3 resulted in the appearance of “core” species, consisting of one GlcNAc linked to asparagine, at 3 positions, namely N88 (14%), N356 (34%), and N616 (25%) (S1 Table, S2 Table “Core” sheet, Fig. 5 in which core is shown in yellow). Concomitantly, there was a partial loss of endo F3-sensitive complex glycans (S1 Table, endo F3-sensitive glycans with scores of 15 and 16 are shaded gray in Fig. 5). However, the quantitative gain of core at each site was not clearly equivalent to the loss of canonical endo F3 substrates. This paradox may be explained by the effects of “NGA” in the glycosidase cocktail, whereby the removal of GlcNAc moieties by N-acetylglucosamidase may reduce glycan score. Thus, for example, a fucosylated trianntennary complex glycans might lose one GlcNAc antenna to become fucosylated biantennary glycan, thereby becoming a potential endo F3 substrate. Further depletion of a GlcNAc antenna from this now fucosylated biantennary glycan would then render it endo F3-resistant once again. Glycan score reduction is perhaps most clearly demonstrated by the appearance of high mannose glycans at N88 and N616 (S1 Table, S2 Table, Fig. 5).

Aside from the 3 positions above, core did not appear at other predominantly complex positions, namely N188, N276, N463, and N611 (S1 Table, S2 Table, Fig. 5). As expected, positions mostly occupied by high mannose glycans (N156, N160, N241, N332, N339, N362, N386 and N448) were unaffected by NGAF3 (S3 Fig). In Fig. 6B, we modeled the core at N88, N356 and N616 in yellow and the unchanged complex glycans at N188, N276, N463 and N611 remained magenta (Fig. 6B). These latter sites may resist NGAF3 due to high glycan density or conformational masking. Although 1% core glycan was detected at position N188 in the NGAF3 sample, similar traces of core at N188, N262, N332, N339, N362 and N448 were also detected in the parent average (S1 Table, S2 Table “Core” sheet).

An apparent increase in high mass complex glycans at N276 in the NGAF3 sample is confounding (Fig. 5). These cannot be products of NGAF3 digestion. Sample variation can be ruled out, as parent VLPs are the source of the NGAF3 sample. Instead, the difference is likely to be experimental: we do not always recover all the structures for every glycan site, which also accounts for modest differences between parent VLP experimental repeats (S1 Table). Lastly, as expected, “N”euraminidase digestion resulted in the loss of sialic acid at all positions occupied by complex glycans, namely N88, N188, N276, N356, N463, N611 and N616 (S2 Table “Sialic acid” sheet), as modeled on Fig. 7A.

**Fig. 7.**
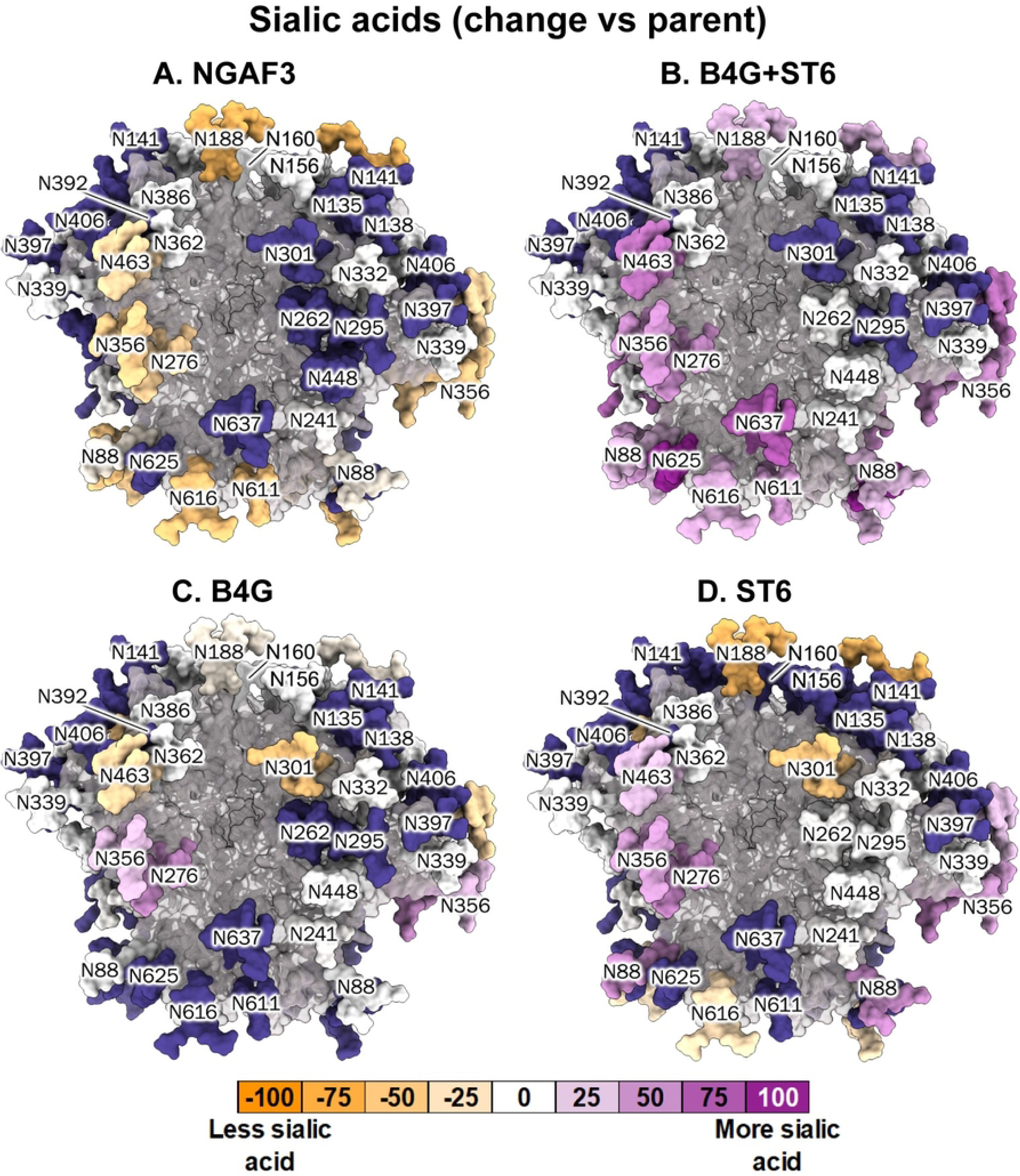
Structural models of sialic acid changes. Related to Fig. 5, S1 Table and S2 Table. Glycans are numbered according to HxB2 strain. The following VLPs are compared to parent VLP average: (A) NGAF3-digested trimer, (B) B4G+ST6 trimer, (C) B4G trimer, and (D) ST6 trimer to show if glycan modification impacted sialylation at each site. Sequons at which sialic acids increase (purple) or decrease (orange) relative to the parent are indicated. Glycans with no data are colored blue.

### ii) B4G+ST6 promotes hybrid glycans in VLP Env

B4G+ST6 induced more pervasive changes at N88, N188, N276, N356, N463, N616, N625 and N637, preventing the maturation of complex glycans beyond hybrid stage (S1 Table, S2 Table “Hybrid” sheet, Fig. 5). Indeed, gp41 glycans N625 and N637 almost completely changed from bulky complex to hybrid. In some cases, hybrid glycan was absent in the parent (N188, N356, N463, N625 and N637); in others, B4G+ST6 amplified pre-existing hybrid glycans (N88 and N616). As expected, predominantly oligomannose glycans N156, N160, N241, N262, N332, N339, N362, N386 and N448 were unaffected (S1 Table, S2 Table “Oligomannose” sheet, S3 Fig). In Fig. 6C, the 7 glycans that substantially became hybrid are shown in brown. In stark contrast, the unchanged N611 glycan remained magenta. Notably, B4G+ST6 and NGAF3 both depleted complex glycans at N88, N356 and N616, resulting in hybrid or core, respectively (Fig. 5, Fig. 6B and 6C). B4G+ST6 may be more penetrative because it works *during* folding rather after folding, as for NGAF3. During synthesis, nascent uncleaved gp160 is relatively flexible, so may be more accessible to B4G and ST6.

### iii) B4G+ST6 increases the prevalence of existing hybrid and sialylated glycans

We next checked the effect of B4G+ST6 on hybrid glycan diversity and abundance. Hybrid glycans were 8.9-fold more abundant on B4G+ST6 than parent VLPs (S4A Fig). All hybrid glycan types were amplified (S3 Table “Summary” sheet). However, only one unique hybrid glycan structure, HexNAc(3)Hex(5)Fuc(1)NeuAc(1), was identified at N276, N356 and N637 (S4B Fig, S4C Fig, S3 Table “B4G+ST6 VLP” sheet). Thus, B4G+ST6 largely increases the abundance of hybrid glycans, but does not introduce many new hybrid structures. This is consistent with the smaller size of B4G gp160/gp120 and the endo H-sensitive gp41 due to hybrid glycans at N616, N625 and N637 (S2B Fig, compare lanes 1 and 2 to lanes 5 and 6).

Sialylated glycans were 1.4-fold more abundant in B4G+ST6 (S5A Fig) but exhibited only 17 different sialylated structures - 2 fold less than parent VLP (S5B Fig). No unique sialic acid structures were observed (S5C Fig). In fact, the parent carries 38 unique sialic acid species that are not present in B4G+ST6 (S5C Fig, S3 Table). Changes in sialic acid abundance per site for B4G+ST6 VLP are modeled in Fig. 7B. Large increases were observed at N88, N188, N276, N356, N463, N616, N625 and N637, all of which also showed increased hybrid glycans (Fig. 6C). Overall, B4G+ST6 modestly increases sialic acid abundance, but reduces their diversity, suggesting a focusing effect, as observed above for hybrid structures.

### iv) Individual effects of B4G and ST6 on VLPs

To check if B4G accounts for hybrid glycans of B4G+ST6 VLPs and for possible B4G-ST6 co-dependency, we examined B4G’s and ST6’s individual effects.

**B4G.** B4G alone promoted hybrid glycans at N88, N188, N276, N356 and N463, as for B4G+ST6 (S1 Table, S2 Table, Fig. 5, Fig 6D). All 4 gp41 glycopeptides were missing in the B4G data set, so we do not know if B4G converts any of these to hybrid glycan (Fig. 5, S2 Table “Hybrid” sheet). Conversely, unlike B4G+ST6, we obtained B4G data at N301. In parent, N301 bore a mixture of oligomannose and complex glycans, whereas in B4G, there was a small complex glycan (96%) and core (4%) (Fig. 5, S1 Table, S2 Table). Thus, like N611 above, N301 is an outlier. Intriguingly, B4G rendered the fully oligomannose N160 glycan partially complex (S1 Table, S2 Table, S3 Fig). Possibly, B4G-mediated glycan thinning at N188 increases local flexibility, allowing the adjacent N160 oligomannose glycan to develop into a complex glycan. The neighboring N156 glycan is also more processed in B4G than both the parent and B4G+ST6 further supporting this possibility.

Compared to the parent, hybrid glycan abundance increased (∼6.4 fold) (S4A Fig), but hybrid diversity decreased (S4B Fig). No new hybrid glycans occurred at any site (S4C Fig). In fact, the parent exhibited 8 unique structures that were not found in B4G (S4C Fig, S3 Table). Thus, like B4G+ST6, B4G increased hybrid glycan abundance but not diversity.

In contrast, B4G reduced sialic acid abundance by 5-fold (S5A Fig). Sialic acid structural diversity was also reduced compared to B4G+ST6 (S5B Fig). Nevertheless, B4G exhibited 2 sialic acid glycans that were absent in the parent, while the parent exhibited 62 sialic acid structures that were absent in B4G (S5C Fig, S4 Table). Changes in sialic acid abundance per site for B4G VLP are modeled (Fig. 7C), showing increases at N276 and N356 and losses at N301 and N463. Collectively, this data reveals a previously unknown effect of B4G in promoting glycan arm extension at the expense of lateral branching. More hybrid glycans were detected by amplifying a few structures, most of which were present in the parent. Conversely sialic acids decreased in frequency and diversity.

**ST6.** ST6 reduced hybrid glycan abundance by 5-fold (S4A Fig), most notably affecting the N616 glycan (S2 Table “Hybrid” sheet, Fig. 5, Fig. 6E). Concomitantly, it reduced hybrid glycan diversity (S4B, S4C Fig). Sialylation increased at some of the same positions as B4G+ST6 (N88, N276, N356 and N463; S2 Table “Sialic acid” sheet, compare Fig. 7B and 7D). However, unlike B4G+ST6, sialic acids were reduced at N188, N301 and N616. Thus, while B4G+ST6 and ST6 can both increase sialic acids, frequently at the same positions, B4G works together with ST6 to promote sialic acids over the entire trimer surface (Fig. 7B and 7D).

Overall, the rank order of modifications for inducing hybrid glycans was B4G+ST6>B4G>parent>ST6 and for inducing sialic acids was B4G+ST6>parent>ST6>B4G, suggesting B4G and ST6 cooperation.

### Impact of glycan density on B4G-mediated hybrid glycans

From the above data, we surmise that B4G promotes hybrid glycans only when space is limited. To test this idea, we examined the effects of B4G on monomeric gp120 (‘gp120’ hereafter) which exhibits reduced glycan density due to its largely glycan-free exposed inner domain. Previously, we found glycan maturation differences between VLP Env parent and gp120 [5, 8]. For unmodified gp120, we calculated the averages of two runs, which showed largely consistent patterns (S1 Table, S2 Table, S6 Fig). In VLPs, sites N156, N160 and N241 were largely populated by high mannose glycans, but were a mixture of high mannose and complex in gp120, presumably due to increased space (S6 Fig). The effects of B4G on gp120 were comparatively mild: we did not see hybrid glycan increases at N88, N276, N356 and N448 (S2 Table “Hybrid” sheet, S6 Fig). However, there were modest increases at N156, N160, N241, N301, N356 and N463 (S6 Fig, S2 Table).

Compared to parent VLPs, hybrid glycans were 21-fold more plentiful on gp120, increasing to a 56-fold with B4G (S4A Fig). Gp120 expressed one novel hybrid structure (S4B Fig, S3 Table) at 3 positions (S4C Fig). There were more novel hybrid glycans on gp120 (regardless of B4G) compared B4G VLPs, where hybrid glycans were homogeneous (S4C Fig).

B4G had mixed effects on gp120 sialic acids, with increases at N88, N160, N241, and N463 and decreases at N156 and N276 (S2 Table “Sialic acid” sheet, S4 Table). Oligomannose increased at N156, N301 and N339 (S6 Fig, S2 Table “Oligomannose” sheet). However, B4G did not impact glycans predominantly occupied by oligomannose (S7 Fig) in either gp120 or VLP formats. Gp120 sialic acid structures were more abundant than on VLPs, and were amplified by B4G (S5A Fig). Some unique sialic acid structures were found on gp120 but not on VLPs (S5B Fig, S4 Table “Unique sialylated glycan” sheet), but B4G reduced this diversity (S5B Fig). At the sequon level, gp120 sialic acid diversity was far higher than VLPs, regardless of B4G (S5C Fig).

Finally, we tested the effect of B4G on MAb VRC01’s N297 glycan. Like gp120, there was little or no change in score (S1 Table). Overall, we conclude that when space is limited, B4G promotes hybrid glycan elongation at the expense of lateral branching.

### B4G+ST6 VLP immunogenicity in rabbits

We showed above that ST6 hypersialylates glycan termini, increasing charge to be more akin to that in PBMC-grown HIV-1 [5]. B4G provides a new way to decrease glycan size and variability, which could allow NAbs to navigate past or engage glycans, potentially increasing vaccine efficacy. This is supported by our previous observation that B4G+ST6 improves sensitivity to PG9, 8ANC195 and 35O22 [5]. The unprecedented role of hybrid glycans in PG9 binding was also shown by another group using glycan arrays [30].

We immunized 3 groups of 4 rabbits (Fig. 8). Groups 1 and 2 received B4G+ST6-modified JR-FL SOS gp160ΔCT E168K VLPs made by co-transfecting MuLV Gag. These VLPs were formulated in AS01B and Adjuplex, respectively. Group 3 received unmodified parent VLPs in Adjuplex. Robust gp120 titers were observed in all rabbits (S8A Fig). Titers were similar in groups 1 and 2, but were 2.4-fold higher in group 3, although this difference was not significant by Kruskal-Wallis test. Gp41 titers were comparable between groups. Unexpectedly, the neutralizing rabbit 613 serum from a previous study [16] reacted several fold more weakly to both gp120 and gp41 compared to the current rabbits. Pooled rabbit sera to gp120 reacted with the cognate immunogen, as expected, but not with gp41. Rabbits immunized with “bald” (no Env) VLPs did not react with either protein. HIV-1+ donor plasmas 1648 and 1686 bound robustly to both gp120 and gp41.

**Fig 8.**
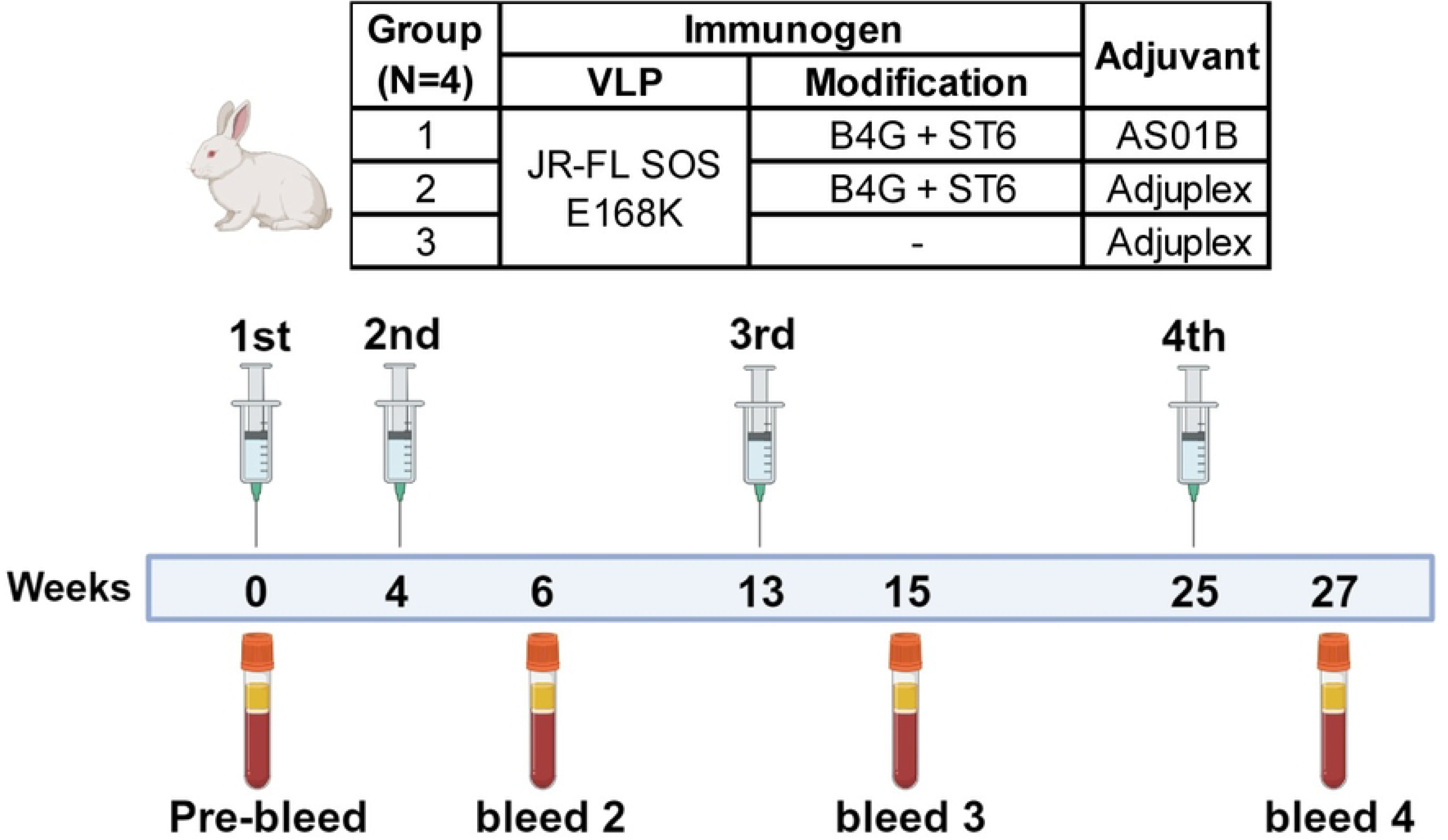
Overview of rabbit immunogenicity studies. Three groups of 4 rabbits were immunized with unmodified Parent VLPs or modified VLPs, as shown. VLP immunogens were produced by transfecting plasmids expressing JR-FL SOS gp160ΔCT E168K and MuLV Gag in 293T cells, with and without B4G and ST6 expression plasmids, and formulated in AS01_B_ or Adjuplex. The week of each immunization is indicated along with the bleeds, taken two week thereafter. Figure was created using BioRender.com.

Next, we evaluated serum reactivity with Env, Gag and host membrane components. VLPs were made using i) mismatched SIV Gag and Env (red dots), ii) Matched MuLV Gag with no Env (purple triangles; i.e., MuLV Gag bald VLPs), and iii) mismatched SIV Gag with no Env (blue squares; i.e., SIV Gag bald VLPs) (S8B Fig). Binding to these VLP variants indicates reactivity to Env, Gag or host protein antigens, as summarized in the S8B Fig. Rabbit sera bound approximately equally to the 3 VLP variants (S8B Fig). Since the blue squares contain neither matched Gag nor Env, this indicates predominant serum binding to host proteins and membranes integral to VLP preparations. making it impossible to unequivocally discern the contributions of anti-Env antibodies. In contrast, rabbits immunized with JR-FL gp120 did not react to either of the “bald” VLPs but reacted to Env VLPs, as expected. Plasmas 1686 and 1648 reacted with the Env VLPs with SIV Gag. 1648 also reacted with “SIV Gag only” bald VLPs, suggesting cross-reactivity of anti-HIV Gag antibodies in this serum with SIV Gag.

NAbs were detected to the JR-FL parent in 5 out of the 12 rabbits: R15 (group 1), R19 (group 2) and R21, R22 and R24 (group 3) (Fig. 9, first column). Parent Env appeared to be superior NAb inducers than B4G+ST6 VLPs both in terms of proportion of responders and average ID50s (424 and 47, respectively). However, the difference was not significant (Kruskal-Wallis, P = 0.0657). As expected, there was also no correlation between gp120 binding titers and NAb ID50s (parent or B4G+ST6 PVs) (r = 0.3138, P = 0.3179). Given that there was only one responder in each of groups 1 and 2, it is unclear if Adjuplex or AS01B offered any advantages.

**Fig 9.**
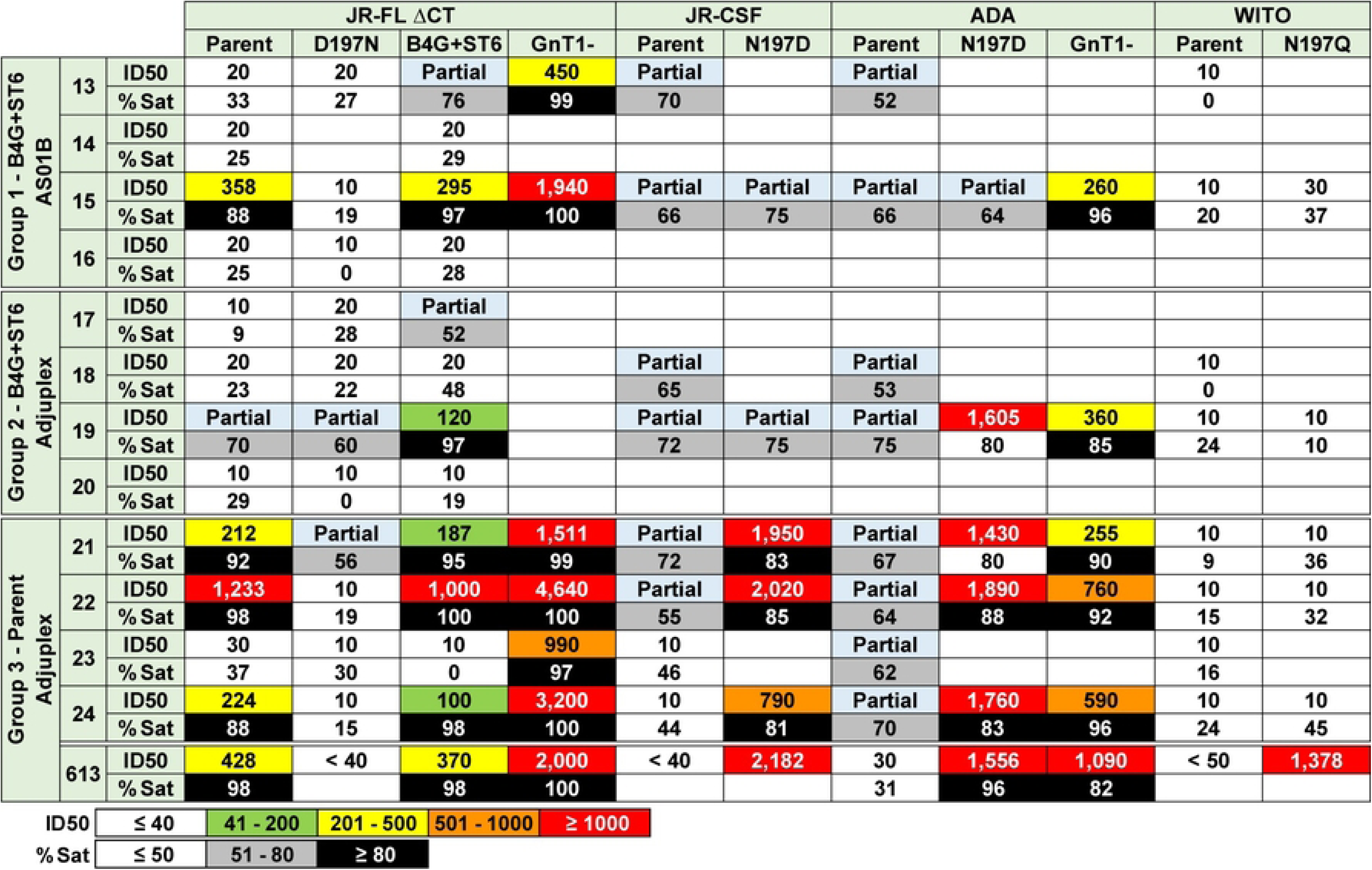
Rabbit serum neutralization of selected subtype B strains. In this heat map, neutralization against selected subtype B strain PVs were recorded as ID50 titers and as percent maximum neutralization saturation (% Sat) at 1:40 serum dilution. Incomplete neutralization is indicated as “Partial” when the % Sat is between 51% and 79%. Progressively warmer-colored cells indicate potent neutralization ID50 titer. % Sat greater than 80% are colored as black, whereas cells with “Partial” neutralization are colored grey.

Groups 1 and 2 showed somewhat improved B4G+ST6 neutralization compared to the parent (Fig. 9, compare columns 1 and 3). A NAb ID50 of 120 was observed for serum R19, whereas it only partially neutralized the parent. Furthermore, partial B4G+ST6 neutralization was detected in R13 and R17 that did not detectably neutralize the parent. This suggests that B4G+ST6 sera cannot navigate the more complex glycans of the parent than they do the cognate B4G+ST6 VLPs. Other rabbits showed serum NAb ID50s to B4G+ST6 PV that were comparable to parent ID50s, regardless of group. Maximum percentage NAb saturation was slightly greater to B4G+ST6 PVs, again regardless of group.

HIV-1+ plasmas neutralized both B4G and B4G+ST6 PVs with improved ID50s compared to the parent PV (∼1.9 and ∼3.3-fold higher) (Fig. 10, S9 Fig). B4G+ST6 improved ID50s was almost significant (Dunn’s test, P = 0.0537), whereas B4G modest ID50s increase was not significant (P = 0.7674). NAb saturation was generally high in most plasmas, but the relatively weak plasmas 1648 and 1652 showed clear increases in saturation to B4G+ST6 (S9 Fig). Overall, broad HIV-1+ plasma NAbs appear to take greater advantage of glycan thinning and α-2,6 sialylated termini driven by B4G+ST6, compared to vaccine sera. However, samples that weakly neutralized the parent were improved for both the HIV-1+ plasmas and rabbit vaccine sera. Rabbit sera with detectable ID50s to parent PV neutralized GnT1-PVs with higher ID50s (between 4 and 33-fold), with improved saturation (Fig. 9). HIV-1+ plasmas also neutralized GnT1-PVs more effectively compared to parent PV (∼10 fold, Dunn’s test P = 0.0004) (Fig. 10, S9 Fig).

**Fig 10.**
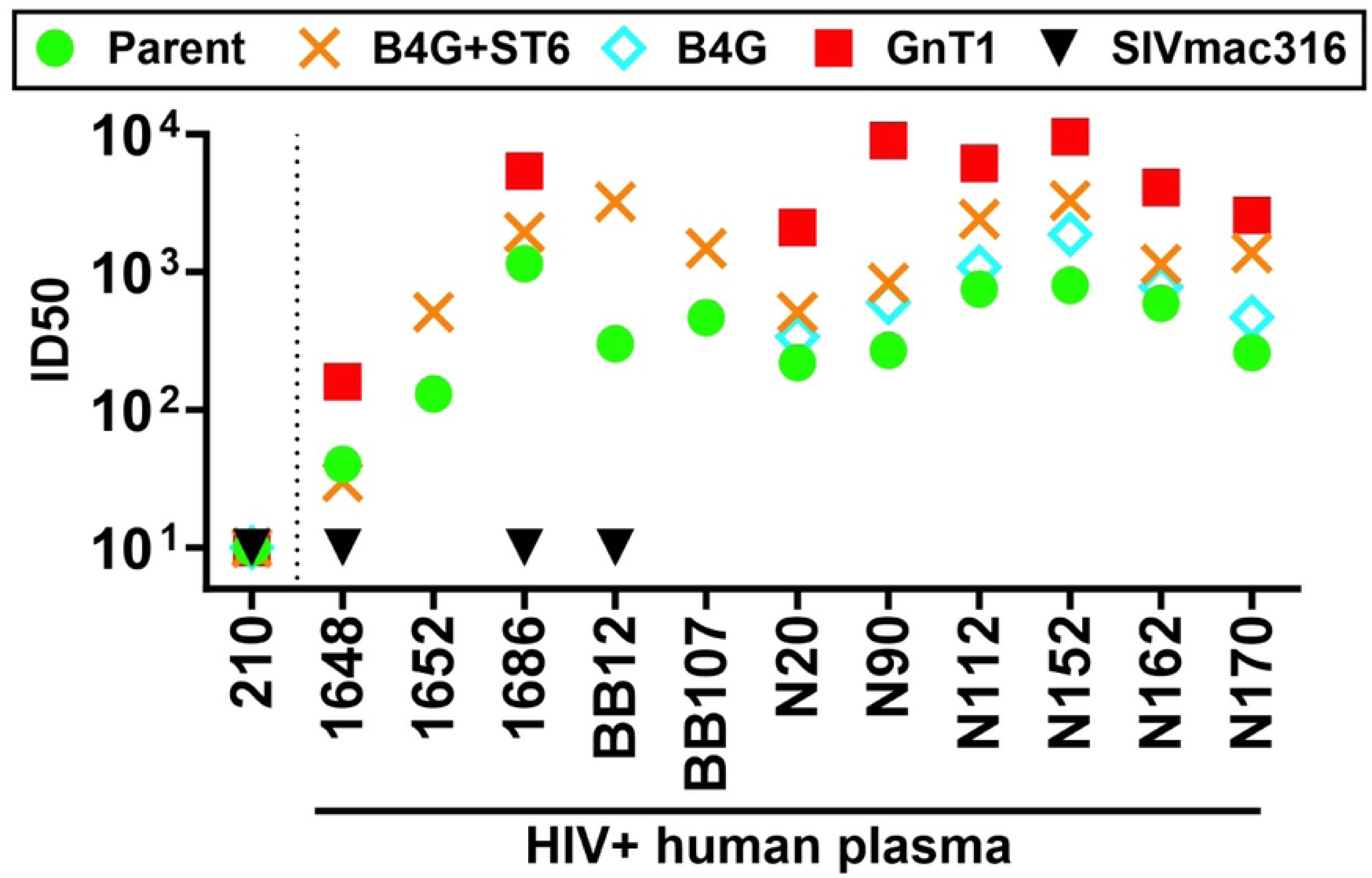
Effect of glycan modifications on HIV-1+ human plasma neutralization. Neutralizing ID50s of a panel of HIV-1+ human plasmas against glycan-modified JR-FL WT gp160ΔCT E168K+N189A or SIVmac316 PVs. 210 is an HIV-negative human plasma.

Collectively, the data suggests that HIV-1+ plasma bNAbs take greater benefit from B4G+ST6 than vaccine sera. Comparing B4G and B4G+ST6 titers (S9 Fig) suggests the presence of bNAbs that contact α-2,6 sialic acids, like PG9 in these plasma, although other specificities like 35O22 are also boosted by these modifications. Conversely, while parent VLP sera cross-neutralized the B4G+ST6 PVs, 3 of 4 neutralizing B4G+ST6 VLP sera (R13, R17 and R19 in Fig. 9) neutralized the cognate B4G+ST6 PVs better than the parent.

### Serum mapping and breadth

Rabbit sera R15 (group 1), R22 and R24 (group 3) did not neutralize the D197N mutant (Fig. 9, S10A Fig), suggesting NAb specificities near the CD4 binding site similar to the 613 rabbit serum that we previously described (Fig. 9). We further investigated using BN-PAGE trimer shifts. R22 (group 3) potently shifted JR-FL SOS E168K trimers (S10B Fig (upper gel), compare lanes 1 and 3), but no shift was observed with D197N trimers (S10B Fig (lower gel)). On the other hand, R13 and R23 poorly shifted both trimers (S10B Fig, compare lanes 1 to 2 and 4 on both gels), consistent with their lack of neutralization (Fig. 9). Overall, the R15, R22 and R24 sera all took advantage of the N197 glycan hole. R19 and R21 sera were less N197-sensitive, evidently targeting another epitope. Titers were insufficiently high to determine if B4G+ST6 NAbs targeted novel epitopes (e.g., R13 in group 1 and R19 in group 2 (Fig. 9)). Rabbit sera with detectable ID50s to JR-FL PVs (R15, R19, R21, R22, R24) also neutralized N197-glycan deficient clones from subtype B strains JR-CSF and ADA, further supporting the glycan hole dependent epitopes of these sera (Fig. 9). This current rabbit sera did not neutralize WITO N197Q, suggesting slightly different specificities from the 613 serum (Fig. 9). Rabbit sera had no detectable neutralization of other subtypes (S11 Fig).

## DISCUSSION

A network of glycans almost completely covers the Env trimer surface, leaving few “protein only” targets for antibodies [4]. Where glycan density is greatest, glycan trimming is very poor, resulting in a conserved “high mannose patch” [32]. Other glycans exhibit variable branching and maturation. For example, N463 is occupied by a range of high mannose and complex glycans, perhaps due to steric competition. This opens up the possibility of epigenetic NAb resistance due to host-encoded glycan variation, a point supported by the increased sensitivity of GnT1-trimers. The present data shows how the glycan network can be molded for vaccine designs.

### How might glycan modifications be best used in successive heterologous vaccine shots?

In recent years, the concept of heterologous prime, boost and polishing immunogens has gained traction based on the rarity of bNAb precursors and the challenges that need overcoming to stimulate and evolve them. In theory, priming elicits a rich diversity of precursor Abs, that may be molded into cross-reactive bNAbs by increased binding constraints and by driving Abs to contact conserved epitopes.

Our data suggests that B4G+ST6 VLPs are best suited as intermediate vaccine boosts, as the increased sialic acid charge appears to diminish NAb priming, consistent with another study [33]. Priming might best be done with heavily glycan-depleted trimers like GnT1-, with increased NAb sensitivity (S9 Fig). Several bNAb precursors bind better to Env trimers produced in GnT1-cells [5, 20, 34, 35]. However, poor expression limits the use of GnT1-cells to produce vaccines. NGAF3 and NGAF123 trimers provide intriguing alternatives. Although we have long considered closed tier 2 trimers to be logical for NAb priming, only 3 of 7 candidate glycans were depleted by NGAF3, achieving slightly improved V2 NAb precursor binding. This might not be enough, and therefore NGAF123 open trimers might therefore be a compromise that maintains fundamental trimer constraints and may prime a richer pool of Abs. It is worth considering that some contemporary candidate priming immunogens, for example fusion peptide conjugates and eOD GT8 do not resemble native trimers [36]. These approaches are, however, designed to be complemented by the use of heterologous closed trimer boosts to filter out NAbs from the priming pool [37]. In contrast, NGAF123 trimers are still functional for infection, supporting their vaccine relevance, and might offer another possible alternative platform from which to boost “on track” NAbs, as compared to subunit primes.

The efficacy of heterologous boosts may in part depend on the “affinity drop” of priming antibodies for the boost, which might need to fall within a certain window to amplify on track NAbs rather than stimulating new ones [38]. B4G could be used to “tune” intermediate vaccine boosts to reduce the affinity drop, thus restimulating a greater number of on track clones. The sera of rabbits 15 and 19 neutralized the parent JR-FL virus with poor saturation compared to B4G+ST6 PV (Fig. 9). In contrast, group 3 sera neutralized parent and B4G+ST6 viruses equally. Thus, B4G-induced bNAbs poorly recognize branched glycans but are no better than “parent” NAbs at B4G virus neutralization. This implies that fully branched glycans are needed in a polishing boost.

Unlike our previous 613 rabbit serum, some of our current rabbit sera partially neutralized clade B strains JR-FL, JR-CSF and ADA with the N197 glycan intact. Neutralization improved when the N197 glycan was absent (N197D) (Fig. 9). This suggests that current rabbit NAbs only partially overlap the N197 glycan, whereas 613 serum NAbs completely blocked by the N197 glycan. A technical mistake marred an attempt to isolate MAbs from these rabbits that could have allowed us to better understand these results.

Like B4G, swainsonine, a drug that blocks MAN2 digestion of the D2/D3 glycan arms, also promotes hybrid glycans at the expense of complex glycans (see Fig. 1 in [5]). Thus, at the crossroads in glycan development where nascent glycans may become hybrid or complex, the balance can be tipped by external “forces” like swainsonine or B4G. Our data suggest that this occurs only when glycans are closely packed. Under normal circumstances, enzymes that create complex glycans appear to be in excess. However, when glycans become too crowded for GnT2-5 to cause further branching, B4G promotes elongation while limiting diversity. By comparison, gp120 glycan score variability (S6 Fig), hybrid glycan diversity (S4C Fig) and sialic acid diversity (S5C Fig) were high and were largely unchanged by B4G.

Paradoxically, B4G+ST6 increased high mannose glycans at N88 and N616. One interpretation is that rapid glycan maturation at one gp160 protomer may prevent maturation at others. Thus, glycosyltransferase overexpression may reduce glycan complexity by out competing other enzymes, resulting in more focused structures that reflect the preferred behavior of these enzymes when effectively isolated from competition.

ST6’s effects are also a reflection of glycan competition (Fig. 7). Excess ST6 accelerates maturation of the most sensitive substrates. The resulting increased charge may force a greater focus on a limited number of structures that minimize repulsion by keeping sialic acid termini separated spatially. The ST6-induced drop in N616 sialylation was coupled with a loss hybrid glycan and gain in oligomannose. N188 and N301 also lost sialic acid (Fig. 7D). Conversely, B4G skews to hybrid glycans at N188 that are apparently able to still accomodate some negative charges. The overall slight loss of sialic acid (S5A Fig) reflects the balance of gains and losses modeled in Fig. 7.

B4G may assist ST6 in two ways that explain the powerful combined effect on the hybrid and sialylated glycan abundance of usually lower diversity. First, B4G helps increase galactose termini, that would allow for sialylation. Second, it thins glycans into monoantennary hybrid glycans that may increase available space for sialylation without lateral repulsion from the sialic acid termini of adjacent glycans. It was previously reported that B4G and ST6 can form homo- or heterodimers regulated by the Golgi environment [39]. Heterodimerization may enhance substrate “channeling” through these two enzymes, which may underlie the observed focusing effect.

High glycan density may explain the less penetrative effects of NGAF3 (N188, N276, N463 and N611 are protected). On the other hand, it could be due to the lack of canonical endo F3 substrates. The fact that NGAF123 evidently removed additional glycans, concomitantly causing a ’globally sensitive’ trimer phenotype, supports the latter explanation. Overall, glycoengineering can reduce glycan complexity or spatial overcrowding which can increase NAb saturation and possibly breadth.

### Why is the N611 glycan uniquely unperturbed by either B4G or NGAF3?

The N611 glycan may be partly protected from endo F3 by being sandwiched between the N616 (and N637) glycan(s) and the membrane at the trimer’s base (Fig. 11). However, the fact that the N611 glycan matures into Env’s heaviest complex glycan implying that it has ample space to do so. Therefore, spatial constraints are lacking for B4G to convert it to hybrid glycan. Nevertheless, the effective removal of sialic acids suggests that the glycan head is accessible (S2 Table “Sialic acid” sheet). Notably, uncleaved I559P Env on membranes exhibits poor N611 glycan maturation (S2 Table “Glycan scores” sheet), suggesting the possibility that proteolytic processing incurs localized refolding that renders the N611 glycan accessible to enzymatic maturation, but remains endo F3-resistant.

**Fig 11.**
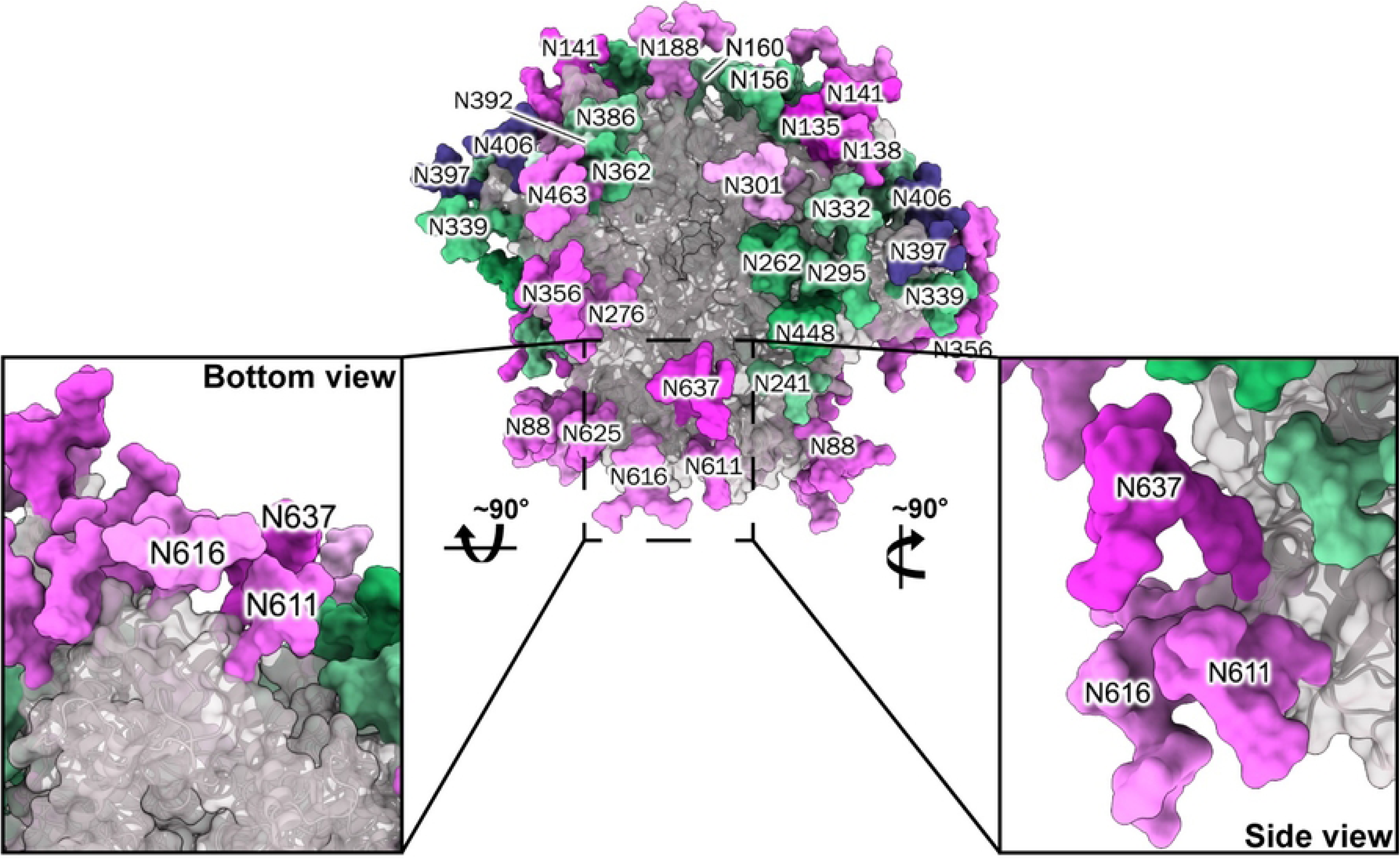
Structure of the HIV-1 Env trimer base. The JR-FL trimer is shown, with the stalk region magnified and rotated to reveal the location of the N611 glycan and its juxtaposition to the neighboring N616 and N637 glycans at the trimer’s base.

### What are the relative effects of glycoengineering and stabilizing mutations on trimer conformation?

LC-MS glycopeptide analysis is a valuable arbiter of Env conformation. We previously investigated the effects of mutations including I559P, NFL and A433P by this method [8]. I559P caused significant changes, consistent with a conformational cost. By comparison, the effects of NGAF3 and B4G are far more subtle and focused (S2 Table “Glycan scores” sheet). Thus, glycan modifications generally do not incur a global Env conformational change, unlike I559P, which widely affects glycan scores and sequon occupation across the trimer surface. However, an exception is NGAF123 which did incur a conformational change, as discussed above. We would like to point out one drawback of the current LC-MS data is that it is done on total samples, so *some* uncleaved gp160 contaminants are present. However, JR-FL gp160 is unusually well processed into gp120/gp41 in 293T cells compared to other strains. Furthermore, we found that trimer-derived purified gp120 exhibits a glycoprofile that largely resembles that of unfractionated VLP Env [8].

Overall, our data further reveals the often huge impact of glycans on bNAb sensitivity. This supports rational vaccine protocols in which glycan clashes are minimized during priming, then incrementally reinstated with greater complexity in boosts to encourage initial glycan-intolerant NAbs to navigate past or engage glycans. In the meantime, LC-MS provides a fascinating simplified window into the molecular interdependencies of Env’s glycans. At present, many of these relationships and patterns are too complex to decipher. However, we envisage that further analysis of Env mutants and glycan variants could be used to understand and correlate glycan maturation dynamics and changes using bioinformatics and structural models.

Our workhorse vaccine platform has been VLPs [16, 40]. However, since VLPs are not readily translatable, they have instead served only as a model for testing membrane trimer vaccine concepts. The development of nucleotide-based vaccines now provides a translatable platform. The modifications we describe here may not be directly adaptable to these new platforms, although modifiers could be used for similar effects.

## MATERIALS AND METHODS

### Plasmids

i. **Env.** Abbreviated Env strain names are given first, with full names and GenBank references in parentheses: JR-FL (JR-FL, AY669728.1), Q23 (Q23.17; AF004885.1), WITO (WITO.33; AY835451.1), c1080 (c1080.c3; JN944660.1), JR-CSF (JR-CSF, AY669726), AC10 (AC10.29; AY835446.1), KER2018 (KER2018.11; AY736810.1), CNE58 (CNE58; HM215421.1), 001428 (001428-2.42; EF117266.1), ADA (AT426119), ZM233 (ZM233.6; DQ388517), 16055 (16055.2; EF117268). Most were in expression plasmids pCI, pCDNA3.1, pCAGGS or pVRC8400. All Envs were truncated at amino acid 709, leaving a 3 amino acid gp41 cytoplasmic tail, termed as gp160ΔCT. This increases native trimer expression and produces pseudoviruses (PV) with similar neutralization sensitivity profiles compared to their full-length gp160 counterparts [8]. Most Env clones are of native amino acid sequence and contain “SOS” mutation (A510C and T506C) to introduce an intermolecular disulfide bond between gp120 and gp41 [41], unless otherwise stated. Additional mutations were introduced to JR-FL Env as follow: E168K+/-N189A mutations knocks in sensitivity to V2 apex bNAbs or A328G mutation converts JR-FL trimers to a tier 1 phenotype (knocks in sensitivity to non-NAbs, due to a conformational effect [8]). All gene synthesis, cloning and mutagenesis were performed by GenScript (USA). Amino acids are numbered according to the canonical HxB2 subtype B reference strain.
ii. **Gag and Rev.** A plasmid expressing Moloney murine leukemia virus (MuLV) Gag (NP_057933.2) was used to produce VLPs [42]. In some instances, simian immunodeficiency virus (SIV) Gag derived from mm251 strain was used (M19499.1). For Env plasmids using native codons, we co-transfected pMV-Rev 0932 that expresses codon-optimized HIV-1 Rev.
iii. **NL4-3.Luc.R-E.** This plasmid is based on HIV-1 proviral clone NL4-3 in which the firefly luciferase gene replaces Nef and does not express Env or Vpr due to frameshift mutations. This plasmid is co-transfected with Env plasmids to produce PVs for neutralization assays.
iv. **Glycosyltransferase plasmids.** Glycosyltransferase plasmids pEE6.4_B4GalT1 (expresses β-1,4 galactosyltransferase and pEE14.4_ST6Gal1 (expresses β-galactoside α-2,6-sialyltransferase were provided by Gillian Dekkers [43].

### MAbs

MAbs were obtained from their producers or the NIH AIDS Reagent Repository. These included PGT145, PG9, PG16, CH01 and CH01 (unmutated common ancestor; UCA), CH04 and CH04 UCA, VRC38.01 and VRC38.01 mHgL, directed to the V2 apex; 14E, 39F, directed to the V3 loop; 2G12 and PGT121, directed to V3-glycan; VRC01, F105 and b12 directed to the CD4 binding site (CD4bs); 35O22 and PGT151, directed to the gp120-gp41 interface and 2F5, 4E10, 10e8, 7B2 and 2.2B directed to gp41.

### HIV-1+ donor plasmas

A collection of HIV-1+ plasmas we and others previously described [44] were obtained from various subtype B- and C-infected donors. Three subtype B plasmas, from United States donors 1648, 1652, 1686, and uninfected donor 210 were obtained from Zeptometrix (Buffalo, NY). Subtype C plasmas (BB12, BB107) were purchased from the South African Blood Bank (Johannesburg). N20, N90, N112, N152, N162 and N170 were obtained from the Vaccine Research Center (VRC) [42].

### Rabbit vaccine sera from previous immunization studies

Previously, we immunized rabbits with either JR-FL Env-VLPs (613), “bald” VLPs devoid of Env (Bald VLP) or gp120 monomer (gp120) [40] (rabbit IDs are shown in parentheses). These sera were used as comparators to the current rabbit immunization study.

### Virus-like particles (VLPs) and pseudovirus (PV) production

For VLP production, Env plasmids were co-transfected with MuLV Gag plasmid and pMV-Rev 0932 in human embryonic kidney 293T (or GnT1-) cells using polyethyleneimine (PEI Max, Polysciences, Inc.). 48h later, supernatants were collected, precleared, filtered, and pelleted at 50,000g. Pellets were washed with PBS, recentrifuged in a microcentrifuge at 15,000rpm, and resuspended at 1,000x the original concentration in PBS.

For PV production, 293T (or GnT1-) cells were co-transfected with Env plasmid and pNL4-3.Luc.R-E using PEI-max. Supernatants were collected 48h later, precleared, filtered and used for neutralization assay.

### Glycoengineered (GE) particles

Glycoengineered (GE) particles (PVs and VLPs) were produced by multiple methods, some of which were described previously [5]. Their effects are summarized in Fig. 1. In brief, GE particles were produced either i) in a N-acetylglucosamine transferase I-deficient GnT1-293S cell line that replaces complex and hybrid glycans with Man5, ii) with pEE14.4_ST6GAL1 plasmid (β-galactoside α-2,6-sialyltransferase 1 – Transfers sialic acid to Galβ1-4GlcNAc structures), iii) with pEE6.4_B4GalT1 (β1,4-galactosyltransferase 1 – transfers of Gal residues to an acceptor GlcNAc by a β1,4-bond), or iv) co-transfection of pEE14.4_ST6GAL1 and pEE6.4_B4GALT1 [5].

### Enzymatic deglycosylation of VLPs

1000x concentrated VLPs were resuspended in 50mM sodium phosphate pH 5.0. To 25µl of 1000x VLP, 1µl (i.e., a molar excess to saturate their effects) of the following glycosidases (used alone or in mixture) was added as follow: Neuraminidases (NA) from *A. ureafaciens* (Sigma) and *C. welchii* (Sigma) capable of removing α-2-3,6,8 sialic acids (SA) (designated as “N”), β(1-4)-Galactosidase (Prozyme) (designated as “G”), β-N-Acetylglucosaminidase (Sigma) with a broad specificity for β-(1,2,3,4,6)-linked GlcNac residues (designated as “A”), Endo F1, F2 (Sigma), derived from *F. meningosepticum)* and Endo F3 (NEB) (designated as F1, F2 and F3, respectively). For eg, “N” VLPs were digested with neuraminidases alone, while “NGAF3” VLPs were digested in a mixture of neuraminidases, galactosidase, acetylglucosaminidase and Endo F3. These mixtures were incubated in PCR tubes at 37°C for 1h. Subsequently, deglycosylated VLPs were washed with 1xPBS and resuspended in fresh 1xPBS.

For endo H digests, VLPs were mixed with 2% SDS and 2-mercaptoethanol and boiled for 5 minutes. Samples were then split in half. To each sample, 2μl of endo H (1000U) (New England Biolabs) or 2μl of PBS (mock) were added and samples were incubated at 37°C for 15 minutes. Samples were then lysed and processed for SDS-PAGE-Western blot.

### Animal immunization

New Zealand white female rabbits were housed and immunized at Covance (Denver, PA), approved by the Association for Assessment and Accreditation of Laboratory Animal Care. Covance animal welfare assurance number is A3850-01. All procedures were approved by IACUC committees at Covance and San Diego Biomedical Research Institute. All animals were fed, housed and handled in strict accordance with the recommendations of the NIH *Guide for the Care and Use of Laboratory Animals* and the Animal Welfare Act. For all procedures, pain and distress were slight and momentary, and did not affect animal health. Discomfort and injury to animals was limited to that which is unavoidable. Analgesics, anesthetics, and tranquilizing drugs were used as necessary by veterinary staff. After the completion of all immunizations and bleeds, rabbits were euthanized by injection of an overdose of anesthesia according to NIH guidelines. Ketamine and xyalzine were administered intramuscularly at 35mg/kg and 5mg/kg, respectively, followed by exsanguination.

Three groups of 4 rabbits were immunized intramuscularly with JR-FL SOS gp160ΔCT E168K VLPs with and without B4G+ST6 modification. VLPs were formulated either in AS01_B_ (Glaxo SmithKline, consisting of liposomes containing deacylated monophosphoryl lipid A and QS-21) or Adjuplex (Advanced BioAdjuvants, consisting of purified lecithin and carbomer homopolymer). VLP doses were determined by analzying their relative antigenicity to JR-FL gp120 by ELISA using various mAbs, as described previously [40]. Immunizations occurred at weeks 0, 4, 13 and 25. Standard serum volumes were drawn on the day of each immunization and two weeks thereafter. Bleeds were heat inactivated at 56°C for 1h prior and stored at -80°C.

### SDS-PAGE-Western blots

VLPs were denatured by heating in 2xLaemmli buffer containing 2-mercaptoethanol (Bio-Rad) for 10 minutes at 95°C, and proteins were resolved in 4–12% Bis-Tris NuPAGE gel (ThermoFisher). Proteins were wet transferred onto a PVDF membrane and blocked in 4% skim milk/PBST. Blots were probed for 1h at room temperature with MAb cocktails in 2% skim milk/PBST, as follows (epitopes in parentheses). Anti-gp120 MAb cocktail: VRC01 and b12 (CD4bs), 39F and 14E (V3 loop), 2G12 and PGT121 (V3 glycan). Anti-gp41 MAb cocktail: 2F5 and 4E10 (membrane proximal ectodomain (MPER)), 7B2 (cluster I), 2.2B (cluster II). After washing, blots were probed with a goat anti-human IgG alkaline phosphatase (AP) conjugate (Accurate Chemicals) at 1:5,000 in 2% skim milk/PBST for 30 minutes at room temperature. Following washing, protein bands on the blots were developed with chromogenic substrate SigmaFast BCIP/NBT (Sigma). All blots were run at least twice to check for reproducibility. Representative data is shown.

### Blue Native (BN) PAGE-Western blots

VLPs were solubilized in 0.12% Triton X-100 in 1mM EDTA. An equal volume of 2x sample buffer (100mM morpholinepropanesulfonic acid (MOPS), 100mM Tris-HCl, pH 7.7, 40% glycerol, and 0.1% Coomassie blue) was added. Samples were spun to remove debris and loaded onto a 4–12% Bis-Tris NuPAGE gel and separated for 3h at 4°C at 100V. Proteins were then transferred to PVDF membrane, de-stained, and blocked in 4% skim milk/PBST. Membrane was probed with a cocktail of mAbs 39F, b12, 14E, PGT121, 2F5 and 4E10, followed by goat anti-human IgG AP conjugate (Accurate Chemicals) and were developed using SigmaFast BCIP/NBT (Sigma).

In BN-PAGE band shifts, VLPs were mixed with sera or mAb for 1h at 37°C, followed by washing with PBS to remove excess ligand and subsequent processing for BN-PAGE as above, using goat anti-human IgG AP conjugate. All blots were run at least twice to check for reproducibility. Representative data is shown.

### Neutralization assays

Neutralization assays were performed as described previously using CF2.CD4.CCR5 cells [5]. Briefly, PV was incubated with graded dilutions of mAbs or sera for 1h at 37°C, then added to CF2Th.CD4.CCR5 cells, plates were spinoculated, and incubated at 37°C for 2h. For SOS PV, following 2h incubation, 5mM dithiothreitol (DTT) was added for 15 minutes to activate infection. All mAb/PV mixture was replaced by fresh media, cultured for 3 days, and luciferase activity was measured using Luciferase Assay System (Promega).

In some cases, 3µg/ml JR-FL gp120 mutant glycoprotein (I423M+N425K+G431E – known as 3mut hereon) was added to test for the ability to interfere with neutralization. These bridging sheet mutations prevent gp120 from binding to target cell receptors [45]. In other cases, excess V3 peptides were added. Eleven residue-overlapping 15-mer consensus clade B V3 peptides were obtained from the NIH AIDS Reagent Program. The peptides used were 8836 (EINCTRPNNNTRKSI), 8837 (TRPNNNTRKSIHIGP), 8838 (NNTRKSIHIGPGRAF), 8839 (KSIHIGPGRAFYTTG) and 8840 (IGPGRAFYTTGEIIG) matches the JR-FL Env sequence were 16used at a final concentration of 3µg/ml each. Neutralization assays were conducted in duplicates and repeated at least three times. Representative data is shown.

### ELISA

ELISAs were performed as described previously [40].

i. **gp120 and gp41 ELISA.** Immulon II ELISA plates were coated overnight at 4°C with recombinant JR-FL gp120 or MN gp41 at 5µg/ml. Following PBST washing and blocking with 4% skim milk/PBST at room temperature, graded dilutions of mAbs or sera were prepared in 2% skim milk/PBST and added to coated glycoproteins for 1h at 37°C.
ii. **VLP ELISA.** Immulon II ELISA plates were coated overnight at 4°C with VLPs at 20x their concentration of transfection supernatants. Following PBS washing and blocking with 4% BSA/PBS/10% FBS at room temperature, titrated mAbs or sera were prepared in 2% BSA/PBS/10% FBS and added to coated VLPs for 1h at 37°C.

Goat anti-human IgG AP or goat anti-rabbit IgG AP conjugate, and PNPP Substrate tablets (ThermoFisher) were used to detect binding. Plates were read at 405nm. Sera dilution resulting in an optical density (OD) of 0.5 (approximately 5 times above background) was recorded as its binding titer. All ELISAs were performed at least twice to check for reproducibility. Representative data is shown.

### Env reduction, alkylation and digestion for mass spectrometry

The following JR-FL SOS gp160ΔCT E168K + N189A VLP samples were subjected to glycopeptide analysis: Untreated, B4GalT1, ST6Gal1, B4GalT1+ST6Gal1 and NGAF3. VLPs were initially buffer exchanged into 50mM Tris HCl, 0.1% DDM (w/w) to disperse lipids. VLPs were then buffer exchanged into 50mM Tris HCl pH 8.0 for subsequent reduction and alkylation.

We also performed glycopeptide analysis on JR-FL gp120 with and without B4GalT1. Prior to reduction and alkylation, JR-FL gp120 was denatured for 1h in 50 mM Tris HCl, pH 8.0 containing 6M urea and 5 mM DTT.

Env proteins were reduced and alkylated with 20mM iodoacetamide (IAA) for 1h in the dark, followed by a 1h incubation with 20mM DTT to eliminate residual IAA. Alkylated Env proteins were buffer exchanged into 50mM Tris HCl, pH 8.0 using Vivaspin columns (3 kDa). Aliquots were digested separately overnight using trypsin and chymotrypsin (Mass Spectrometry Grade, Promega). The next day, peptides were dried and extracted using C18 Zip-tip (Merck Millipore).

### Liquid chromatography-mass spectrometry (LC-MS) glycopeptide analysis

Peptides were dried and resuspended in 0.1% formic acid, and analyzed by nanoLC-ESI MS with an Ultimate 3000 HPLC (Thermo Fisher Scientific) system coupled to an Orbitrap Eclipse mass spectrometer (Thermo Fisher Scientific) using stepped higher energy collision-induced dissociation (HCD) fragmentation. Peptides were separated using an EasySpray PepMap RSLC C18 column (75μm × 75cm). A trapping column (PepMap 100 C18 3μM 75μM x 2cm) was used in line with the LC prior to separation with the analytical column. LC conditions were as follows: 280min linear gradient consisting of 4–32% acetonitrile (ACN) in 0.1% formic acid over 260 minutes, followed by 20 minutes of alternating 76% ACN in 0.1% formic acid and 4% ACN in 0.1% formic acid to ensure all the sample elutes from the column. The flow rate was set to 300nL/min. The spray voltage was set to 2.7 kV and the temperature of the heated capillary was set to 40°C. The ion transfer tube temperature was set to 275°C. The scan range was 375−1500 m/z. Stepped HCD collision energy was set to 15%, 25% and 45% and the MS2 for each energy was combined. Precursor and fragment detection were performed with an Orbitrap at a resolution MS1 = 120,000, MS2 = 30,000. The AGC target for MS1 was set to standard and injection time set to auto which involves the system setting the two parameters to maximize sensitivity while maintaining cycle time.

### Site-specific glycan analysis

Glycopeptide fragmentation data were extracted from the raw file using Byos (Version 3.5; Protein Metrics Inc.). Glycopeptides were evaluated in reference to UniProtKB <u>Q6BC19</u> (ectodomain of JR-FL gp160ΔCT). All samples carry mutations SOS (A501C, T605C) and E168K and N189A, and others, as denoted in the.txt files found in the MassIVE database (submission in process). Data were evaluated manually for each glycopeptide. A peptide was scored as true-positive when the correct b and y fragment ions were observed, along with oxonium ions corresponding to the glycan identified. The MS data was searched using the Protein Metrics “N-glycan 309 mammalian no sodium” library with sulfated glycans added manually. All charge states for a single glycopeptide were summed. The precursor mass tolerance was set at 4 ppm and 10 ppm for fragments. A 1% false discovery rate (FDR) was applied. Glycans were categorized according to the composition detected.

HexNAc(2)Hex(10+) was defined as M9Glc, HexNAc(2)Hex(9−3) was classified as M9 to M3. Any of these structures containing a fucose were categorized as FM (fucosylated mannose). HexNAc(3)Hex(5−6)X was classified as Hybrid with HexNAc(3)Hex(5–6)Fuc(1)X classified as Fhybrid. Complex glycans were classified according to the number of HexNAc subunits and the presence or absence of fucosylation. As this fragmentation method does not provide linkage information, compositional isomers are grouped. For example, a triantennary glycan contains HexNAc(5) but so does a biantennary glycans with a bisect. Core glycans refer to truncated structures smaller than M3. M9Glc-M4 were classified as oligomannose glycans. Glycans containing at least one sialic acid were categorized as NeuAc and at least one fucose residue in the “fucose” category.

Glycans were categorized into I.D.s ranging from 1 (M9Glc) to 19 (HexNAc(6+)(F)(x)). These values were multiplied by the percentage of the corresponding glycan divided by the total glycan percentage excluding unoccupied and core glycans to give a score corresponding to the most prevalent glycan at a given site. Arithmetic score changes were then calculated from the subtraction of these scores from one sample against others, as specified.

### Statistical analysis

All graphs were generated and analyzed using Prism (version 10.1.1).

## ACKNOWLEDGEMENTS

This work was supported by NIH grants AI00790 and AI93278 (JMB) and International AIDS Vaccine Initiative (IAVI) through grant INV-008352/OPP1153692 funded by the Bill and Melinda Gates Foundation (MC).

## SUPPLEMENTARY FIGURE LEGENDS

**S1 Fig. Neutralization interference by V3 peptides and gp120 3mut.**

Neutralization sensitivity of the glycan variants using the same mAbs in Fig. 3 was further evaluated in the presence of V3 peptides or JR-FL gp120 3mut (I423M+N425K+G431E) using CF2.CD4.CCR5 cell targets. Error bar represents the standard deviation of the mean.

**S2 Fig. Western blot analysis of glycan modified Env.**

(A) BN-PAGE Western blot analysis of JRFL SOS gp160ΔCT E168K VLP parent and glycan modified VLPs, as indicated. Blots were probed with anti-gp120 and anti-gp41 mAb cocktails. Trimer and monomer bands are indicated by colored dots. (B) The same VLPs were boiled in SDS/DTT, treated with or without endo H, then analyzed in duplicate SDS-PAGE Western blots that were probed with either anti-gp120 mAb cocktail (top) or anti-gp41 mAb cocktail (bottom). Env species are indicated by colored dots.

**S3 Fig. The effect of glycan modification on the oligomannose cluster on Env VLPs.**

Related to Fig. 5, S1 Table and S2 Table. Glycans are numbered according to HxB2 strain. Distribution of glycan species is shown at sites predomninantly occupied by oligomannose glycans. The matched Parent VLP is compared against B4G+ST6, B4G, ST6 and NGAF3 VLP. For N386 (underlined), we show the average Parent VLP data because data was missing in the matched Parent VLP. Glycans at position where no glycopeptide was detected are indicated as “No data”.

**S4 Fig. The effect of glycan modification on hybrid glycan frequency and diversity.**

Related to S3 Table. (A) The total eXtracted Ion Chromatogram (XIC) area of all hybrid glycans was calculated for each sample and tabulated in S3 Table. The XIC area for Parent VLP was an average of three independent glycopeptide MS, while the XIC area for gp120 WT was an average of two independent glycopeptide MS. The heat map shows the fold change of XIC area of all hybrid glycans between samples. (B) Bar chart showing the number of different hybrid glycans found in parent VLP, glycan modified VLPs and monomeric gp120 with and without B4G. The number of different hybrid glycan for each sample is indicated above each bar. (C) The heat map showing the presence of unique hybrid glycan (in all positions) for a particular sample in relation to its comparator. Of interest, one unique hybrid glycan species was found in B4G+ST6 VLP, gp120 WT and gp120 B4G samples, but was absent in Parent VLP (See orange highlight in S3 Table “B4G+ST6 VLP”, “gp120 WT” and “gp120 B4G” sheets).

**S5 Fig. Effect of glycan modification on sialylated glycan frequency and diversity.**

Related to S4 Table. (A) This heat map shows the fold change of XIC area of all sialylated glycans between samples. (B) Bar chart showing the number of different sialylated glycans (hybrid and complex) found in parent VLP, glycan modified VLPs, and monomeric gp120 with and without B4G. Number of different sialylated glycan for each sample is indicated above each bar. (C) The heat map showing the presence of unique sialylated glycan (in all positions) for a particular sample relative to its comparator. A list of these unique sialylated glycans present in B4G VLP, ST6 VLP, gp120 WT and gp120 B4G, whereas they are absent in Parent VLP, can be found in S4 Table “Unique sialylated glycan” sheet.

**S6 Fig. Glycan distribution changes at key sites of Env on gp120 and VLPs.**

Related to S1 Table and S2 Table. The histograms show different glycan distributions at these sequons in monomeric gp120 and VLPs with and without B4G. Positions where no glycopeptide was detected are indicated as “No data”.

**S7 Fig. Oligomannose distribution at key sites of monomeric gp120 and VLPs.**

Related to S1 Table, S2 Table and S6 Fig. The histograms show the oligomannose distribution at these sequons in monomeric gp120 and VLPs with and without B4G. Positions where no glycopeptide was detected are indicated as “No data”.

**S8 Fig. Binding profiles of rabbit vaccine sera and human plasmas.**

(A) Recombinant gp120, gp41 and (B) VLP ELISA binding titer of immunized rabbits from the current study (R13 – R24), previous studies (613, gp120 and Bald VLP), and HIV-/HIV-1+ human plasmas. The latter rabbit sera and HIV-1+ human plasmas were included as controls. Env VLP was prepared using JR-FL Env and SIV Gag (mismatching rabbit immunogen that was prepared with MuLV Gag – red circles), MuLV “bald” VLPs (made using MuLV Gag, matching rabbit immunogens – purple triangles) and SIV “bald” VLPs (made using SIV gag, mismatching rabbit immunogens – blue squares). The purposes of using these VLPs are as follow: (1) SIV “bald” VLP measures only host protein responses; (2) MuLV “bald” Gag VLPs measures both gag and host protein responses; (3) Env VLPs with SIV Gag to measure anti-Env-and anti-host protein responses, if any. Each point was representative of data derived from at least 2 repeats.

**S9 Fig. HIV-1+ plasma neutralization of selected subtype B strains.**

Related to Fig. 10. In this heat map, neutralization is presented in ID50 titers and as % maximum neutralization saturation (% Sat) at plasma dilution of 1:40. Progressively warmer-colored cells indicate potent neutralization ID50 titer. % Sat greater than 80% are colored as black, whereas cells with “Partial” neutralization are colored grey.

**S10 Fig. Rabbit vaccine sera mapping.**

(A) Neutralization of selected rabbit vaccine sera against JR-FL WT gp160ΔCT E168K PV with and without D197N mutation on CF2.CD4.CCR5 cells. Error bar represents standard deviation of the mean. (B) Rabbit vaccine sera were mixed with VLPs at 37°C for 1h, washed, lysed and analyzed by BN-PAGE Western blot, probing with anti-gp120+anti-gp41 mAb cocktail. The upper gel was JR-FL SOS gp160ΔCT E168K, while the lower gel was JR-FL SOS gp160ΔCT E168K+D197N. Env species are indicated with colored dots.

**S11 Fig. Limited rabbit serum neutralization of other HIV-1 subtypes.**

Related to Fig. 9. Rabbit vaccine sera were checked for neutralizing activity against AC10 (subtype B) and non-subtype B isolates. N.D. = not done.

## SUPPLEMENTARY TABLE LEGENDS

**S1 Table. Raw glycopeptide data.**

**S2 Table. Glycopeptide components and analysis.**

**S3 Table. Hybrid glycan quantification and frequency.**

**S4 Table. Sialylated glycan quantification and frequency.**

